# Chromatin structure and context-dependent sequence features control prime editing efficiency

**DOI:** 10.1101/2023.04.15.536944

**Authors:** Somang Kim, Jimmy B. Yuan, Wendy S. Woods, Destry A. Newton, Pablo Perez-Pinera, Jun S. Song

## Abstract

Prime editor (PE) is a highly versatile CRISPR-Cas9 genome editing technique. The current constructs, however, have variable efficiency and may require laborious experimental optimization. This study presents statistical models for learning the salient epigenomic and sequence features of target sites modulating the editing efficiency and provides guidelines for designing optimal PEs. We found that both regional constitutive heterochromatin and local nucleosome occlusion of target sites impede editing, while position-specific G/C nucleotides in the primer binding site (PBS) and reverse transcription (RT) template regions of PE guide-RNA (pegRNA) yield high editing efficiency, especially for short PBS designs. The presence of G/C nucleotides was most critical immediately 5’ to the protospacer adjacent motif (PAM) site for all designs. The effects of different last templated nucleotides were quantified and seen to depend on both PBS and RT template lengths. Our models found AGG to be the preferred PAM and detected a guanine nucleotide four bases downstream of PAM to facilitate editing, suggesting a hitherto-unrecognized interaction with Cas9. A neural network interpretation method based on nonextensive statistical mechanics further revealed multi-nucleotide preferences, indicating dependency among several bases across pegRNA. Our work clarifies previous conflicting observations and uncovers context-dependent features important for optimizing PE designs.

## INTRODUCTION

One of the most powerful gene editing tools available today is the complex of clustered regularly interspaced short palindromic repeats (CRISPR) with CRISPR-associated protein 9 (Cas9) (1–4). In nature, CRISPR-Cas9 is found in bacteria as a natural defense mechanism of excising foreign DNA in the CRISPR DNA regions. In a rapid succession of development, researchers have repurposed and further engineered this process as a laboratory tool for editing the genome of a variety of cell types across species, including human diseased cells (5–7). Several different techniques have been developed to date to improve the CRISPR-Cas9 system to be more programmable and suitable for *in-vivo* editing, while reducing off-targets and unintended mutations (8–13). In particular, prime editors (PEs) represent the latest state-of-the-art, highly versatile tool (14). The biomolecular architecture of PEs consists of a partially inactivated Cas9 fused to a reverse transcriptase and a customizable prime editing guide RNA (pegRNA), which contains a scaffolding sequence bound by Cas9. A PE targets the desired edit locus via the combination of two main processes: the complementary pairing of a ∼20 nucleotide (nt) guide sequence at the 5’ end of pegRNA to the non-edited DNA strand and the recognition of a short protospacer adjacent motif (PAM) on the edited strand by Cas9. The modified Cas9 includes a domain fused to a reverse transcriptase (RT) and a nickase domain that nicks only the edited DNA strand, 3 bases upstream of the PAM sequence. The 3’ end of pegRNA hybridizes to a region in the edited strand and acts as a primer for the ensuing reverse transcription. Starting at the nick site, the RT reverse transcribes part of the pegRNA 3’ extension that is immediately upstream of the primer binding site (PBS) region. This region is denoted as the RT template and contains the complementary RNA template for the desired DNA edit sequence. After nicking the edited strand and reverse transcribing the templated DNA, two single-stranded DNA flaps are formed: the 3’ flap created via reverse transcription contains the edit of interest, and the 5’ flap, created via the Cas9 nick, does not contain the edit. Successful edits are completed upon the action of 5’-flap-specific endonucleases, such as FEN1 (15).

There are two main advantages of PEs over other genome editors. First, compared to the CRISPR-Cas9 endonuclease construct, PEs are less likely to produce unintended nearby insertions and deletions, as they avoid DNA double strand breaks (14,16). Second, unlike base editors that can currently introduce only C > T or A > G conversions, PEs can produce all 12 base changes as well as small insertions and deletions (11,14). However, a notable difficulty in using PEs stems from the complication that the flexible pegRNA design has several adjustable parameters, yielding varying degrees of editing efficiency, and from the fact that there is currently a dearth of reliable computational models capable of *a priori* predicting these differences in efficiencies.

Anzalone *et al*. originally introduced three variants of PE, denoted as PE1, PE2, and PE3, where PE1 used a reverse transcriptase derived from Moloney murine leukemia (MMLV RT), PE2 used the MMLV RT from PE1 with an additional 5 point mutations, and PE3 used the Cas9 nickase in PE2 to perform non-concurrent nicks on both strands and thus perform non-concurrent edits on both strands to avoid creating double-stranded breaks (14). To date, numerous engineering approaches have been made to improve the editing efficiency of PE, such as mutating the PE components or co-expressing additional components together with PE, resulting in multiple PE variants (17–21). There have also been additional improvements in PE design by either altering the pegRNA outside the regions directly hybridizing with the target site (22–24), or by impeding the mismatch repair mechanism (20).

Despite the rapid experimental progress, accurately predicting the editing efficiency of PEs at previously untested genomic loci remains a major challenge. The prediction problem is complicated by the fact that PE efficiency may depend on numerous factors, such as the sequence content and chromatin accessibility of the targeted locus, the lengths of the PBS and RT regions on pegRNA, and the intended editing type. Notably, Kim *et al.* recently embarked on the difficult task of probing PE2 efficiency in human cells by using lentiviral plasmid libraries to screen the efficiency of ∼48,000 pegRNA designs on ∼2,000 integrated target sequences (25); analyzing the resulting data using a deep learning model, they predicted the measured PE2 efficiency based on an extensive set of variables including the target sequence, GC counts, melting temperature, minimum self-folding energy, and the DeepSpCas9 score of Cas9 nuclease activity (26). Although the authors highlighted the most relevant features using the Tree SHAP approach, some predictive variables, such as GC content and melting temperature, were highly correlated and might have confounded the interpretation. In addition, their analysis focused on common features shared across varying lengths of PBS and RT template, rather than features distinguishing different designs. Furthermore, the experimental method measuring the editing efficiencies mostly assayed exogenously integrated sequences rather than endogenous sequences. The specific locations of the integration sites were thus not known; as a result, Kim *et al.*’s data and models did not incorporate epigenetic information in predicting PE2 efficiency.

This study presents statistical models that systematically examine the effects of both epigenomic and sequence-dependent features on PE efficiencies by analyzing data from both existing publications and additional in-house experiments (Supplementary Method S1). We consider only PE2 editors, which have by far the most amount of publicly available data. We describe preferred target features for each pair of PBS length (PBSL) and RT template length (RTTL), revealing specific nucleotide effects that depend on these lengths and resolving in the process discrepancies in the literature regarding the role of certain nucleotides. Our models, utilizing only a small number of parameters compared to previous approaches (25), capture both marginal and joint effects of nucleotides across pegRNA and provide practical guidelines for choosing optimal PE2 designs.

## MATERIALS AND METHODS

### Cell culture and transfection

The cell line HEK293T was obtained from the American Tissue Collection Center (ATCC) and was maintained in DMEM supplemented with 10% fetal bovine serum and 1% penicillin/streptomycin at 37°C with 5% CO2. HEK293T cells were transfected in 24-well plates with Lipofectamine 2000 (Invitrogen) following the manufacturer’s instructions. The amount of DNA used for lipofection was 1 μg per well. Transfection efficiency was routinely higher than 90% as determined by fluorescent microscopy or flow cytometry following delivery of a control GFP expression plasmid.

### Plasmids and Cloning

The plasmids encoding PE2 (Plasmid#132775) as well as the plasmid for expressing the pegRNA (Plasmid #132777) were obtained from Addgene. PegRNAs were cloned into the plasmid backbone with paired oligonucleotides (IDT, Supplementary Table S1) as previously described (14). Briefly, the oligonucleotides used to create the guide sequences were hybridized, phosphorylated and cloned into the sgRNA vector using BsaI-HFv2 (NEB), in a reaction that included T4-PNK (NEB) and T4 DNA Ligase (NEB).

### Next Generation Sequencing (NGS)

DNA amplicons for NGS were generated by PCR using KAPA HiFi HotStart (Roche), according to manufacturer’s instructions, using primers with overhangs compatible with Nextera XT indexing (IDT, Supplementary Table S2). Following validation of the quality of PCR products by gel electrophoresis, the PCR products were isolated using an AMPure XP PCR purification beads (Beckman Coulter). Indexed amplicons were then generated using a Nextera XT DNA Library Prep Kit (Illumina) quantitated, and pooled. Libraries were sequenced with a MiSeq Nano Flow Cell for 251 cycles from each end of the fragment using a MiSeq Reagent Kit v2 (500-cycles). FASTQ files were created and demultiplexed using bcl2fastq v2.17.1.14 Conversion Software (Illumina). Deep sequencing was performed by the Roy J. Carver Biotechnology Center at the University of Illinois at Urbana-Champaign.

### Quantification of editing efficiency

Editing efficiency data for PE2 at endogenously edited sites in Anzalone *et al.* were obtained from the Sequence Read Archive (https://www.ncbi.nlm.nih.gov/sra/) under the accession code PRJNA565979. In-house experimental sequencing data for 25 target sites can be found in SRA under the accession code PRJNA949853. Raw sequences were aligned to the human genome (GRCh38) using the Bowtie package (version 2.4.1), with a maximum fragment length of 500 base pairs (bp) for paired-end sequences (27). Poorly aligned sequences were filtered using the Samtools package (version 1.7) with options -h -F 4 -q 10 (28). Editing efficiency for each technical replicate was calculated by dividing the number of reads that contain the edit of interest, but were otherwise perfectly aligned, by the number of perfectly aligned reads containing the wild-type sequence. For paired-end sequencing data, half of the number of reads found in strand pairs were excluded from both the mutant and wild-type sequences to prevent double-counting of reads. The final editing percentage for each target loci was determined by averaging over the editing percentages in all technical replicates.

### Assessment of differential enrichment of histone modifications

Genomic locations of H3K9me3 and H3K27me3 histone modifications were determined from the aggregate of chromatin immuno-precipitation sequencing (ChIP-seq) datasets in HEK293 cells from the Encyclopedia of DNA Elements (ENCODE) Consortium (29,30), with the accession codes having the prefix “ENC,” and the Gene Expression Omnibus (GEO) (31), with the accession codes having the prefix “GSM.” For H3K9me3, the IP sequencing data used were GSM4301086 (32), the combination of ENCFF002AAX and ENCFF002AAZ as replicates, and the combination of GSM3452796 and GSM3452797 also as replicates (33). The corresponding inputs were GSM4301092, ENCFF000WXY, and GSM4445881 (34). Raw reads were aligned to GRCh38 with Bowtie (version 2.4.1). For H3K27me3, the IP sequencing data were GSM3907592 (34), GSM4301076 (32), GSM4586041 (35), and GSM4859391 (36); the corresponding inputs were GSM3907592, GSM4301081, GSM4586041, and GSM4859385. To determine the consensus genomic locations of H3K9me3 enrichment relative to input, we first constructed 6 row-vectors from the *ℓ*_1_-normalized density of reads, partitioned into 1kb genomic bins, with the 3 IP data and 3 input data treated as separate samples. We performed singular value decomposition (SVD) on the resulting matrix of the 6 row-vectors (Supplementary Method S2). A similar analysis was performed to detect the consensus genomic locations of H3K27me3 enrichment relative to input. SVD and further analysis of variance (ANOVA) were performed using the Decomposition and Classification of Epigenomic Tensors (DeCET) package (37).

### Calculation of RNA-DNA hybridization energy

RNA-DNA hybridization energies were obtained by computing the difference in length-normalized Gibbs free energy (Δ*G*°) at 37°C between a paired RNA-DNA oligomer and two unpaired oligonucleotides. The Δ*G*° values of paired oligonucleotides were computed by adding up all the Δ*G*° values of the dinucleotide components of the oligonucleotides, in addition to a helix initiation term that accounts for forming the first base pair in the double helix. The Δ*G*° values of all paired dinucleotides and initiation terms were obtained from Sugimoto *et al.* (38).

### Summary statistic of nucleosome occupancy signal

Nucleosome occupancy in the lymphoblastoid cell line GM12878 were determined from MNase-sequencing bigwig tracks from the ENCODE portal under the accession code ENCFFOOOVME (30). Average nucleosome occupancy in particular genomic range was calculated by determining the sum of MNase-seq signals at every genomic coordinate in the protospacer region, normalized by the length of the protospacer.

### Off-target determination

PE off-targets were computed by aligning the protospacer sequence to regions in the hg38 genome that are also upstream of an NGG PAM site. A genomic locus was considered to be an off-target if its aligned sequence matched the reference protospacer sequence up to three mismatches outside the PAM site and if there were no mismatches in the GG dinucleotide of the PAM motif. These off-targets were determined by using the Cas-OFFInder package (39).

### Elastic net model: linear regression with combined *ℓ*_1_ and *ℓ*_2_ penalities

Linear regression with combined *ℓ*_1_ and *ℓ*_2_ penalties were computed in R (version 4.2.1) using the function glmnet.cv in the glmnet (version 4.1-4) library (40). Given a set of *N* sequences, sequence *S_i_* of length *L* and and edit percentage *E*(*S_i_*) was modeled according to the equation

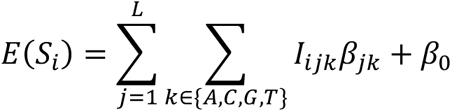

where *I_ijk_* is the indicator variable for the presence of A,C,G or T at the *j*th position in the *i*th sequence, *β_jk_* are the regression coefficients for the indicator variables, and *β*_0_ is the constant intercept. The vector of regression coefficients is estimated as

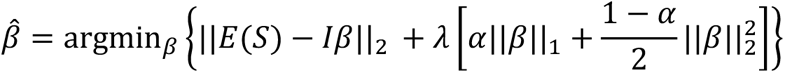

where *E*(*S*) is the vector of edit percentages, and *λ* and *α* are parameters to be tuned via cross-validation. After performing 10-fold cross-validation with a default value of *α* = 0.5, the value of *λ* was tuned to minimize the mean cross-validated error across all 10 folds of the observed edit percentage compared to the predicted edit percentage of the withheld validation dataset.

### Ordinary least-square (OLS) linear regression model for stepwise difference in editing efficiency of consecutive RTTLs using last two templated nucleotides

It was previously observed that having G as the last templated nucleotide at RTTL= *i* tended to decrease the editing efficiency compared to the shorter pegRNA design with RTTL= *i* − 1 (14). In the absence of G, we noticed that the nucleotide A had a similar affect as G. We thus hypothesized that the stepwise differences in editing efficiency between two consecutive RTTLs can be predicted by the last templated nucleotides at those RTTLs. To confirm our hypothesis, we built a linear regression model represented by the equation

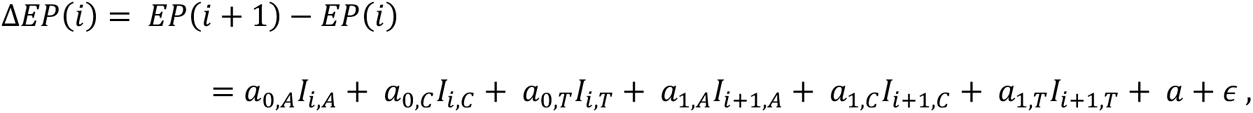

where *EP*(*i*) is the edit percentage at RTTL= *i* in the range [10nt, 20nt] for each target locus, *I_i,N_* is the indicator variable for nucleotide *N* being the *i* th templated nucleotide, the *a*’s are the regression coefficients, and *ϵ* is an error term.

Kim *et al*.’s data contained editing efficiencies for PBSL=13 at only nonconsecutive RTTLs (RTTL=10, 12, 15, and 20); thus, the OLS was applied on the sum of stepwise differences in three ranges of RTTL: (1) ∑_10≤*i*<12_Δ*EP*(*i*), (2) ∑_12≤*i*<15_ Δ*EP*(*i*), and (3) ∑_15≤*i*<20_ Δ*EP*(*i*). To assess the performance of the OLS model, 8-fold or 10-fold cross validation was performed using Anzalone *et al.*’s data or Kim *et al.*’s data, respectively. The target loci were partitioned into eight (or ten) sets and OLS was trained on the Δ*EP*(*i*) (or ∑*_i_* Δ*EP*(*i*)) at the target loci in the union of seven (or nine) sets (training set) where 10 ≤ *i* < 20. The OLS was validated on the remaining held-out set (test set). This process was repeated eight (or ten) times permuting the test set.

### Structure of the deep neural network (DNN) for predicting prime editor efficiency

We trained a DNN model to predict the edit percentages of all pegRNA designs from Kim *et al.* (25) and thereby learn the salient sequence features influencing prime editor efficiency. The inputs to the DNN were 47×5 matrices, where the dimension 47 was chosen to accommodate the length of the “wide-target sequences” in Supplementary Table 4 of *Kim et al.* The first four columns of the input matrix were one-hot encoding of the edited strand sequence at locations from −21 to +26 relative to the nick site. To account for the fact that a given site could be targeted by different pegRNA designs of varying lengths, the fifth column of the input matrix was used to one-hot encode for the coverage of the corresponding locations by each input pegRNA.

To capture position-specific effects of sequence features relative to the nick site, we used independent filters for different positions in the target sequence, instead of using convolutional filters with shared weights. The input sequences were thus divided into all possible overlapping 8-mers, resulting in 40 8-mers for a sequence of length 47nt. Each 8-mer was passed through 10 filters of kernel size 8 × 5, and the resulting output was then flattened to a vector. The flattened output was processed through a fully-connected layer of 10 neurons. The output of the fully-connected layer was then passed through a single neuron whose output was the prediction of the edit percentage of the pegRNA design at the corresponding target site. All layers of the DNN used a rectified linear unit (ReLu) activation function. To prevent overfitting, one dropout layer, where 20% of layer outputs were set to 0, was applied after the input layer and another dropout layer was applied after the filter layer. The DNN was constructed and trained using the Python package Keras version 2.2.0.

### DNN training and interpretation

The full dataset from *Kim et al.* was divided into 80% training, 10% validation, and 10% test sets, grouping together pegRNA designs with the same target site but with different PBSLs or RTTLs into a common set. Since the edit percentage of approximately 60% of the pegRNA designs was less than 10%, we assigned training weights to each pegRNA design in the training set in order to balance the dataset and prevent the DNN from learning biased features from over-represented lowly edited sites. For this purpose, the training set was partitioned according to their observed edit percentages, with a bin size of 1%. A pegRNA design, indexed by *i*, was given a weight

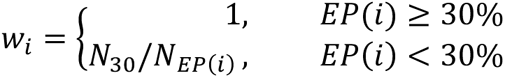

where *N_EP_* is the number of pegRNAs in the partition corresponding to the range [*int*(*EP*)%, *int*(*EP* + 1)%). The above weights were applied to the loss function during DNN training. Training was terminated when there was no improvement in the weighted mean-squared error loss between the predicted log(*EP*) and the observed log(*EP*) for the validation set after 10 consecutive epochs. The DNN model at the epoch with the lowest validation loss was considered for further analysis. The Pearson correlation was determined between the predicted and observed edit percentages in the test set. The DNN was trained using the stochastic gradient descent optimizer with Nesterov momentum of 0.9, initial learning rate of 10^-3^, and decay factor of 10^-6^.

To extract sequence features learned by the DNN, we used a modified version of the simulated annealing (SA) algorithm to determine the optimal sequences that maximize the output of the DNN (41,42); we then used the maxEnt algorithm to identify salient positions and nucleotide preferences in the optimal sequences (43) (Supplementary Methods S3 and S4).

## RESULTS

### Constitutive heterochromatin impedes PE2 efficiency

In order for a PE to be able to edit its target site, the spacer region of pegRNA first must access and hybridize to the complementary DNA sequence at the targeted locus. We thus hypothesized that target sites residing in heterochromatin, a condensed state of highly packaged DNA, may be blocked from access by PEs and thereby have low editing efficiency. Consistent with our reasoning, it has been previously observed that chromatin structure can interfere with the CRISPR-Cas9 endonuclease activity (44–47). We thus investigated the effect of heterochromatin on PE efficiency using genome editing data from Anzalone *et al*. (14), Kim *et al*. (25), and additional in-house validation experiments. To determine heterochromatin locations in HEK293 cells, which were used in all three studies, we integrated several publicly available H3K9me3 and H3K27me3 ChIP-seq datasets as a proxy for indicating closed chromatin regions (Methods). After obtaining consensus regions of enrichment for each of these histone modifications using singular value decomposition of the joint IP and input data matrices (Methods; Supplementary Method S2; Supplementary Fig. S1), we computed the genomic distances from each target site to the nearest H3K9me3 and H3K27me3 modified regions and used these distances as features for predicting editing efficiency. When considering edit percentage as a function of the distance from the protospacer to the nearest H3K9me3 modified region, we observed that the edit percentages at sites close to H3K9me3 were approximately 0%, whereas higher edit percentages were possible farther away (Supplementary Fig. S2A).

This observation motivated us to search for binary classification of target sites as being either weakly or strongly editable, choosing an efficiency threshold of 1% to binarize the data. We then trained a multivariate logistic regression model to classify all 32 endogenous target sites in Kim *et al*., using the distances to H3K9me3 and H3K27me3 modified regions as features (Methods; Supplementary Method S2). The probability threshold separating the binary editing categories was set to maximize the finite positive likelihood ratio of the true positive rate to the false positive rate. Performing the *t*-test for each regression coefficient and determining the bivariate and univariate logistic regression decision boundary showed the H3K27me3 variable to be insignificant (*p* = 2.52 × 10^-1^; Supplementary Table S3; Supplementary Fig. S2B). After removing the insignificant feature and retraining the model, the probability threshold that maximized the finite positive likelihood ratio corresponded to a distance threshold of 26kb for the nearest H3K9me3 peak from the protospacer (Fig. 1A). We observed a substantial increase in PE efficiency at the target sites located at least ∼10 kb away from H3K9me3 modification, suggesting that chromatin assumed an open conformation around this distance threshold. Similarly, it was reported that transcription of genes started to increase ∼10 kb away from H3K9me3 peaks (48). Using the 26kb distance threshold, the accuracy of the classifier on the test set consisting of the data from Anzalone *et al.* and additional in-house validation experiments was 75% with an area under receiver operator characteristic curve (AUROC) of 0.90 (Fig. 1B,C). Our findings thus support that heterochromatin may be sufficient to block PEs from targeting; however, heterochromatin is likely not necessary to hinder PE targeting efficiency, and other factors may also contribute to low edit rate, as discussed below.

**Figure 1.**
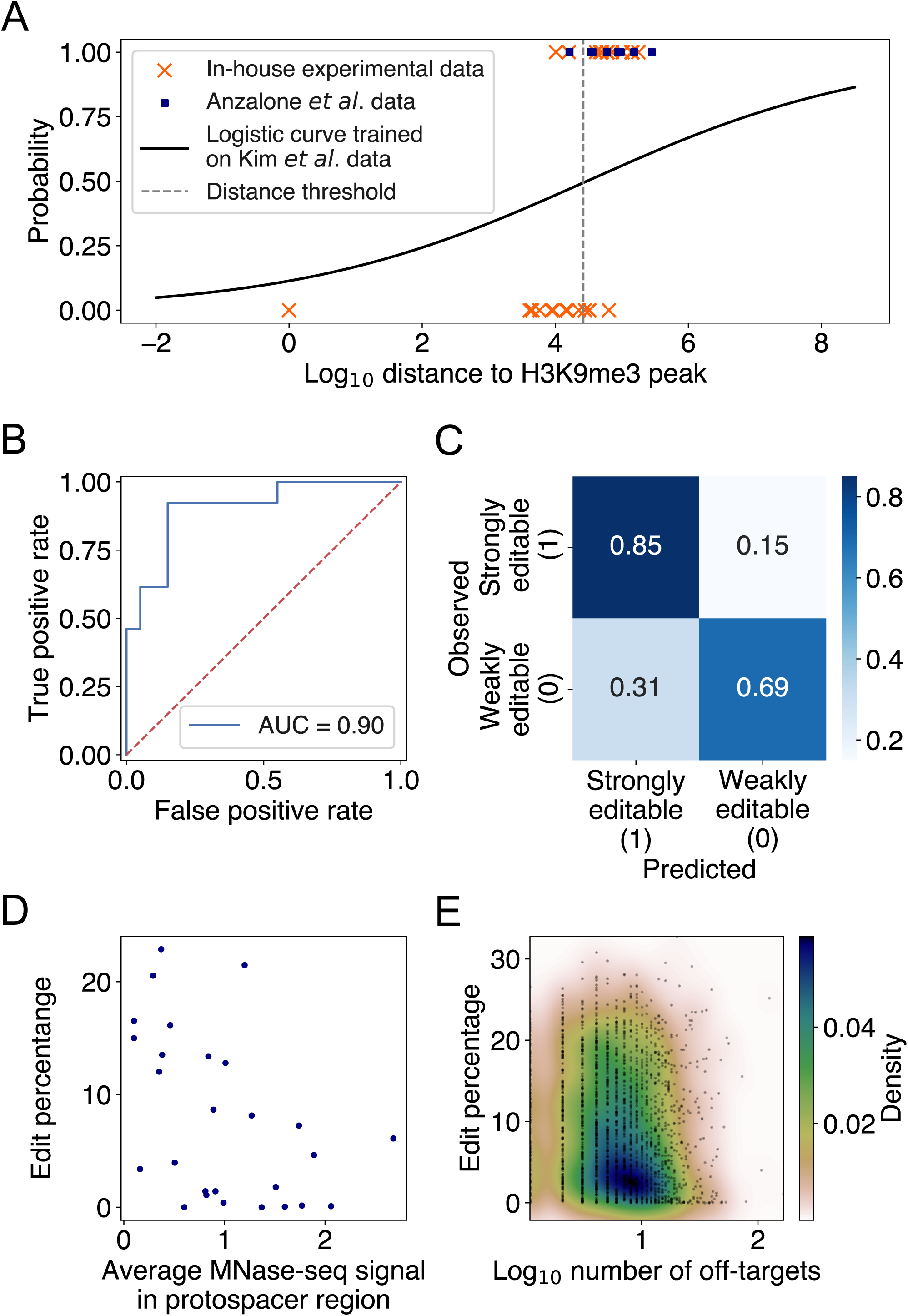
Constitutive heterochromatin near target sites impedes prime editing. (A) Scatter plot of binary classes of editability as a function of log_10_ distance from target site to the nearest H3K9me3 peak (Methods, Supplementary Method S2). The distance threshold (dashed grey line) is at ∼26kb. The logistic regression was trained on 32 endogenous targets from Kim *et al.* and tested on editing data from Anzalone *et al.* and our in-house experiments. (B) Receiver operator characteristic curve (ROC) for the logistic regression evaluated on the test set, with an area under ROC (AUROC) of 0.90. (C) Confusion matrix for the logistic regression predictions on the test set. Overall accuracy was 75%. (D) Scatter plot of edit percentage as a function of MNase-seq signal averaged over the protospacer region (Pearson *r* = −0.49, *p* = 8.72 × 10^-3^) using the endogenous target sites from Kim *et al*. and Anzalone *et al*. that are at least 26kb away from H3K9me3 peaks. Edit percentage for each target locus with different pegRNA designs was determined by averaging the edit percentages of all pegRNA designs sharing the same protospacer. (E) Scatter plot of edit percentages averaged over all integrated target sites sharing the same protospacer as a function of log-transformed number of potential off-targets. The colors represents the density of data points in the scatter plot.

### Nucleosome occlusion and pegRNA off-targets may decrease prime editing efficiency

Having confirmed that high-order chromatin accessibility affects PE efficiency, we next investigated the effect of local chromatin structure on prime editing. It has been previously reported both *in vitro* and *in vivo* that nucleosomes inhibit the efficiency of CRISPR-Cas9 endonuclease, suggesting that histone proteins may block the target DNA access to Cas9 (46,49). Given the shared components between the CRISPR-Cas9 and PE constructs, we thus sought to test whether nucleosome positioning at the target sites might also inhibit PE editing efficiencies. Since there were no publicly available MNase-seq datasets for HEK293 and most nucleosome positioning was shown to possess some degree of consistency across different cell lines (50) and partially exhibit intrinsic DNA sequence preferences (51–54), we used the available nucleosome occupancy data in the lymphoblastoid cell line GM12878 (Methods). We found a significant negative correlation between PE edit percentage and nucleosome occupancy in the endogenous target sites of Kim *et al*. and Anzalone *et al*. (Pearson *r* = −0.46, *p* = 1.36 × 10^-2^; Fig. 1D; Methods). While target sites in nucleosomal DNA tended to have low edit rates, target sites in nucleosome-free regions did not necessarily have high edit rates. Similar to the effect of heterochromatin, our findings thus support that nucleosomes are sufficient, but not necessary, to partially block PEs from targeting, as other factors may also contribute to low edit rate, as shown in the following sections.

Given that the PBS and spacer regions of pegRNA must hybridize to the target DNA sequence in order for editing to occur, with the spacer being the longer of the two sequences, we examined the off-target effect of having several genomic loci complementary to the spacer region on editing efficiency. We found only a weak but statistically significant negative correlation between the number of spacer off-targets and the edit rate averaged over all pegRNAs with the same spacer (Spearman *ρ* = −0.11, *p* = 7 × 10^-7^; Fig. 1E; Methods). This result indicates that the presence of off-targets does not substantially affect the editing efficiency at the intended target and is consistent with the previous finding for single guide RNAs (sgRNA) of CRISPR-Cas9 (55).

### Last two templated nucleotides embody stepwise differences in PE2 efficiency

Our logistic regression model above has revealed that target sites close to heterochromatin are unlikely to be editable. Once a target site away from heterochromatin is selected, an important next step is to optimize the pegRNA design by adjusting the lengths of PBS and RT template. It was previously reported that altering the PBS length (PBSL) and RT template length (RTTL) could have a drastic effect on PE2 efficiency even when targeting the same site for the same edit and that having G as the last templated nucleotide tended to decrease the PE2 efficiency (14). We further observed in Anzalone *et al*.’s data that: 1) at the *EMX1* locus containing no G’s in the RT region (Anzalone *et al.* Fig. 2B), having A as the last templated nucleotide consistently resulted in lower edit rates, suggesting that other nucleotides apart from G might also modulate the editing efficiency; 2) at the *FANCF* locus (Anzalone *et al.* Fig. 2B), even though the last templated nucleotide was G at RTTLs of both 10 and 18 nts, the edit rate for 18 nts was much higher than that for 10 nts, suggesting that the presence of G alone could not explain the variability of edit rates as a function of RT template sequence composition. These findings motivated us to use the sequence content in target region as features in predicting the PE2 efficiency as PBSL and RTTL were varied. We thus developed an ordinary least squares (OLS) linear regression model to predict stepwise differences in edit percentages between two adjacent RTTLs, using the last templated nucleotides as features (Methods).

**Figure 2.**
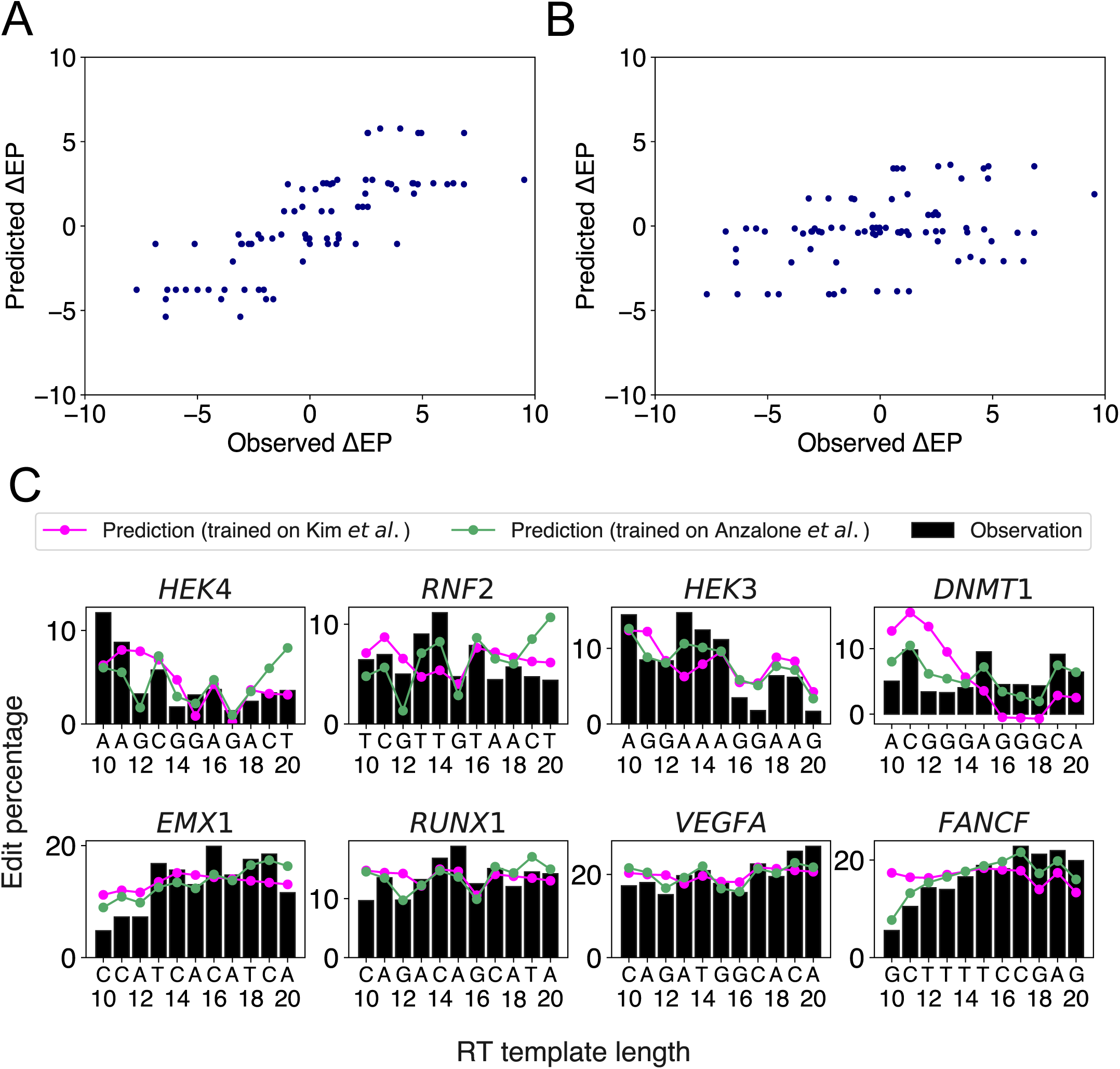
OLS linear regression using the last two templated nucleotides robustly predicts stepwise differences in edit percentage between two consecutive RTTLs. (A) Scatter plot of the predicted versus observed differences in edit percentage when the OLS linear regression model was trained and tested on Anzalone *et al.*’s data. (B) Scatter plot of the predicted versus observed differences in edit percentage when trained on the integrated sites from Kim *et al.*’s data and tested on Anzalone *et al*.’s data. (C) Predicted and observed edit percentages as a function of RTTL for 8 different target sites from Anzalone *et al*. For each target, the absolute predicted edit percentage at RTTL=10 was set such that the average of predicted edit percentages across RTTLs matches the corresponding average of observed edit percentages.

Anzalone *et al*. generated edit percentages (defined as the fractions of total reads with correct edits) of PE2 at eight different gene loci using pegRNAs with varying ranges of RTTL for each locus, with the largest overlap of lengths among the designs being between 10 and 20 nts. We thus chose this range [10nt, 20nt] to train a OLS linear regression model for predicting the stepwise differences, Δ*EP*(*i*) = *EP*(*i* + 1) − *EP*(*i*), in edit percentages between *i* + 1 and *i* RTTL (Fig. 2A, Pearson *r* = 0.79; Methods). Eight-fold cross validation of holding out and testing on each gene locus supported its robustness, with the root mean square error (RMSE) of the test set being mostly similar to that of the training set (∼2.3%) (Supplementary Table S4). Our model learned the effect of the last templated nucleotide on Δ*EP*(*i*) to be in increasing order of G, A, C, T for the (*i* + 1)^th^ templated nucleotide and decreasing order of G, A, C, T for the *i*^th^ templated nucleotide (Supplementary Table S5). The relative ordering pf G<A<C<T resembles the order of ionization energy of the nucleotides (56), but clarifying the connection would require further experimental and theoretical investigation.

Having confirmed that our OLS linear regression model performed well on the eight target sites from Anzalone *et al*., we further investigated whether a similar approach could be generalized to Kim *et al*.’s data (25), which measured editing efficiencies at thousands of target sites using high-throughput sequencing. The PBSL was set to 13 nts, to be consistent with the range used in Anzalone *et al*. Since Kim *et al*. measured the edit percentages only for RTTL=10, 12, 15, and 20, we could not directly apply our stepwise approach on these data. We thus divided the data for different RTTLs into three ranges and trained an OLS linear regression model to predict consecutive stepwise differences in edit percentages for each range separately (Methods): 1) RTTL from 10 to 12; 2) RTTL from 12 to 15; and 3) RTTL from 15 to 20 (Supplementary Fig. S3). Ten-fold cross validation verified the robustness of our model within each range (5.1% RMSE; average of Pearson correlation across folds = 0.33 for RTTL in the range [10,12] and [15,20], and 0.25 for RTTL of [12,15]; Supplementary Table S6). However, the effect sizes of the four nucleotides differed somewhat across the three ranges (Supplementary Tables S7-S9). When RTTL is in the range [15,20], the effect on Δ*ER*(*i*) was in increasing order of G, A, T, C for the (*i* + 1)^th^ templated nucleotide, and decreasing order of G, A, C, T for the *i* ^th^ templated nucleotide (Supplementary Table S9), similar to the aforementioned results for Anzalone e*t al*.’s data. RTTL in the [12,15] range also showed the pattern that G as the (*i* + 1)^th^ templated nucleotide was associated with the smallest Δ*EP*(*i*) and G as the *i*^th^ templated nucleotide was associated with the largest Δ*EP*(*i*) compared to other nucleotides at these respective positions (Supplementary Table S8). This pattern was reversed for the RTTL in the [10,12] range, where G as the *i* ^th^ templated nucleotide was associated with the smallest Δ*EP*(*i*) compared to other nucleotides at this position (Supplementary Table S7).

Testing the regression model trained on Kim *et al.*’s data on Anzalone *et al*.’s data in the corresponding RTTL ranges (Supplementary Tables S7-S9), the predicted Δ*EP*(*i*) still significantly correlated with the observed values, albeit to a lesser extent than the model directly trained on Anzalone *et al*.’s data (Pearson *r* = 0.37, *p* = 7.13 × 10^-4^; Fig. 2A,B). Repeating the same analysis in individual RTTL ranges revealed that the predicted Δ*EP*(*i*) was significantly correlated with the observed values only in 15-20 RTTL range (Pearson *r* = 0.25, *p* = 0.34 in 10-12 RTTL range; Pearson *r* = 0.05, *p* = 0.81 in 12-15 RTTL range; Pearson *r* = 0.62, *p* = 2.12 × 10^-5^ in 15-20 RTTL range). We also attempted to predict the trend of absolute edit percentages in Anzalone *et al*.’s data as a function of RTTL (Fig. 2C). Overall, the model trained on Anzalone *et al*.’s data was able to reproduce the observed trend that the edit percentage decreased whenever the last templated nucleotide was G (green line in Fig. 2C). The models trained on Kim *et al.*’s data were able to capture the observed trend in longer RTTL, but not so well in shorter RTTL (magenta line in Fig. 2C), consistent with the fact that our predicted Δ*EP*(*i*) was more accurate in the long RTTL range [15,20].

There were some discrepancies between the two independent data sets regarding the regression coefficients trained on shorter RTTL designs. For example, Anzalone *et al*. observed a decrease in editing efficiency whenever the last templated nucleotide was G throughout all RTTL. By contrast, Kim *et al*. observed that, on average, editing efficiency was highest when the last templated nucleotide was G at RTTL=10 and 12, while it was lowest for the last templated G only at RTTL=20 (Kim *et al*. Fig. 2F). Further investigation is needed to understand why G has an opposite effect on editing efficiency at short RTTLs and why this phenomenon is not universal across independent data sets. We shall revisit the position-specific effects of G on editing efficiency in subsequent sections.

### Elastic net regression accurately predicts editing efficiency using the sequence content of target DNA and flanking regions

The OLS linear regression analysis yielded insight into preferred RTTLs of pegRNAs for a given target site, but it could not reveal the optimal target, given multiple PAM candidates in the region of interest. We therefore built an elastic net model (40), a linear regression model with both *ℓ*_1_ shrinkage to impose sparsity of features and *ℓ*_2_ shrinkage to reduce overfitting, for each combination of PBSL and RTTL, separately (PBSL=7, 9, 11, 13, 15, 17 and RTTL=10, 12, 15, 20) (25) and predicted the absolute editing efficiency of a target site using only the sequence information of PBS, RT template and its flanking regions (Methods). The predictive variables were the indicator variables encoding the four nucleotides at each position of the 47 bp-long target site sequence, except for the fixed GG dinucleotide in the NGG PAM (Methods). Since this dataset did not contain information about where the target sites were integrated in the genome, features involving relative distance to epigenomic modifications were not included in our model. Our approach was much simpler and more interpretable than the previously reported model pooling together different PBSL and RTTL designs and using 1,766 features (25), many of which might have been highly correlated and redundant.

The elastic net predictions significantly correlated with the observed edit percentages in each pair of PBSL and RTTL (Pearson *r* > 0.60, *p* < 10^-150^; Supplementary Table S10). Fig. 3A shows the regression coefficients for PBSL=13 and RTTL=15 (other combinations of PBSL and RTTL in Supplementary Fig. S4). In general, the higher the GC content in the PBS region, the more likely the target site was to have high editing efficiency (Supplementary Fig. S5), consistent with the previous report (25). However, the extent of this preference depended on the PBSL; that is, the high-performance designs with shorter PBSL required higher GC content in the PBS region than those with longer PBSL (Supplementary Fig. S4). Considering that G:C bonds are stronger than A:T bonds, this result suggested that a short PBS lacking a sufficient number of G/C nucleotides may not stably hybridize to the 3’ end of pegRNA, thereby yielding low editing efficiency. To further investigate this idea, we computed the length-normalized RNA-DNA hybridization energy between the pegRNA PBS region and the corresponding DNA and found a significant negative correlation between PBS-DNA hybridization energy and edit percentage for all combinations of PBSLs and RTTLs (Fig. 3B,C; Supplementary Fig. S6; Methods). Additionally, we confirmed again that the shorter the PBSL, the larger the magnitude of the anti-correlation between editing efficiency and PBS-DNA hybridization energy (Fig. 3B vs. Fig. 3C, Supplementary Fig. S6).

**Figure 3.**
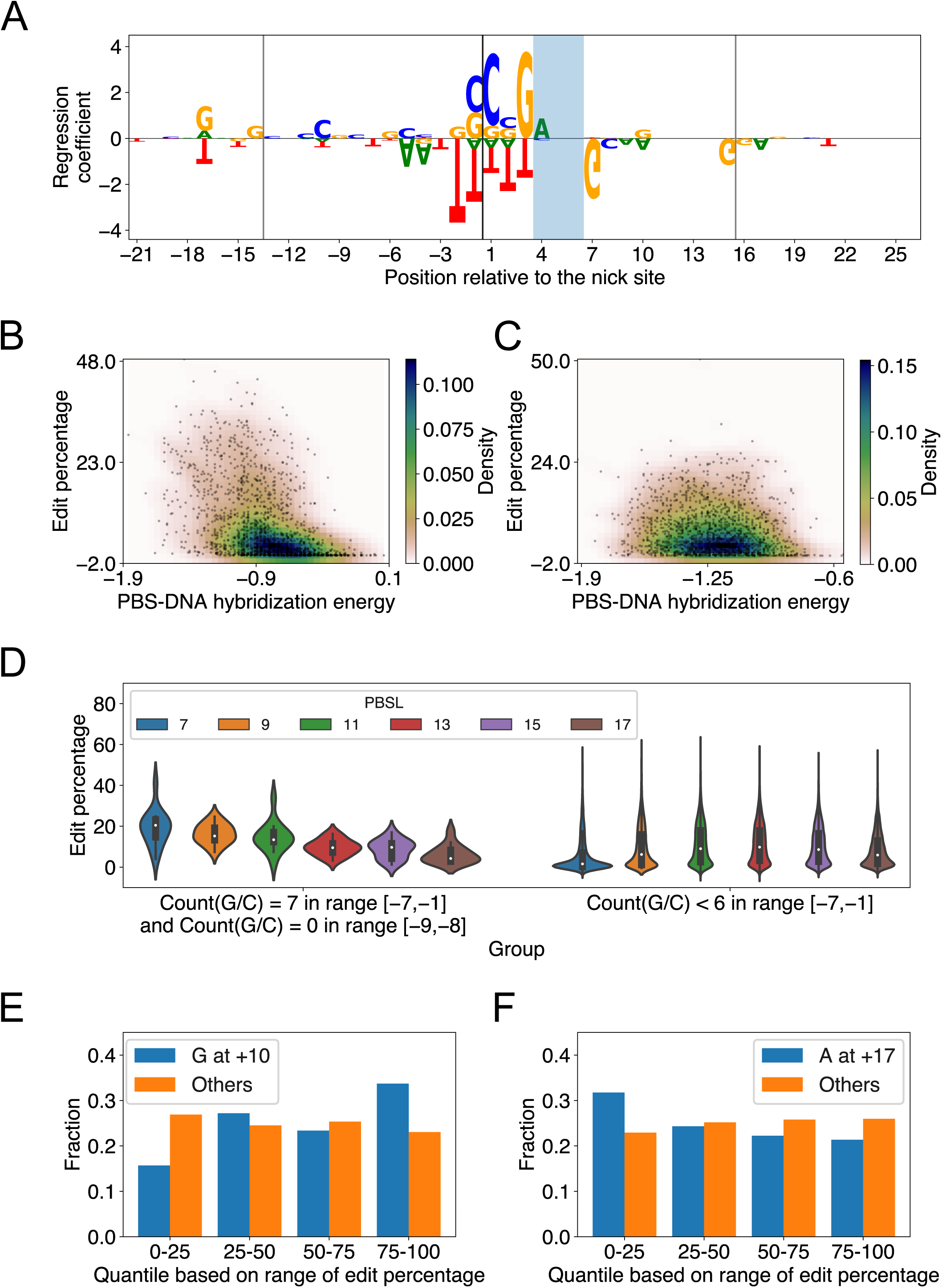
Elastic net regression model learns sequence features important for predicting PE2 efficiency. (A) Elastic net regression coefficients for all edited-strand nucleotides in the range [-21,+26] relative to the nick site for PBSL=13 and RTTL=15. The model was trained on the editing data for integrated target sites from Kim *et al*. The nucleotide heights represent the absolute value of the corresponding regression coefficients. The vertical lines denote the borders of the PBS and RT template regions, and the PAM site is shaded. (B) Scatter plot of edit percentage versus PBS-DNA hybridization energy for PBSL=7 and RTTL=10, where the colors represent the density of data points. Pearson *r* = −0.45, *p* = 1.1 × 10^-78^. (C) Same as in (B), but for PBSL=17 and RTTL=20. Pearson *r* = −0.13, *p* = 2.8 × 10^-7^. (D) Violin plot of edit percentages for each PBSL in two groups of target sites, divided based on the GC composition of the 9 bp-long sequence upstream of the nick site. The first group contains target sites that have only G or C in the range [-7,-1] and only A or T at positions −9 and −8. The second group contains target sites that have less than six G or C nucleotides in the range [-7, −1]. The white dots represent the median edit percentage, and the thick blank lines represent the portion of the data between the first and third quartiles of edit percentage. (E) Bar plot denoting the fraction of target sites in each quantile of edit percentages. Blue (or orange) bars are for target sites with (or without) G at position +10 for PBSL=13 and RTTL=15. (F) Same as in (E), but for A at position +17.

The regression coefficients also showed that the presence of G as the last templated nucleotide tended to increase editing efficiency when RTTL=10 regardless of the PBSL, or when RTTL=12 and PBSL ≤ 9 (Fig. 3A, Supplementary Fig. S4). This finding recapitulated the observation made by Kim *et al*. that the editing efficiency was on average highest when the last templated nucleotide was G for RTTL=10 or 12 (25); however, our analysis clearly highlighted that this effect depended on the PBSL being sufficiently short for the case RTTL=12. By contrast, when PBSL ≥ 11 and RTTL ≥ 15, the presence of G as the last templated nucleotide decreased editing efficiency (Fig. 3A, Supplementary Fig. S4), consistent with the findings of our OLS model above and Anzalone *et al*. (14); for shorter PBSL, however, the last templated nucleotide had only a minor effect on editing efficiency, and the GC content in the PBS region instead had a pronounced effect.

The large effect size of the GC content in PBS region for short PBSL designs, together with the accompanying reduction in the negative effect of the last templated nucleotide, suggested that optimal choices of pegRNA for target loci containing high GC content in the PBS region would involve short PBSL. Supporting this idea, we observed that when the pegRNA contained only G/C in the range [-7,-1] and A/T at −9 and −8, the edit percentage was on average highest for PBSL=7 (Fig. 3D; highest median edit percentage of 20.5% when PBSL=7); here, we considered A/T at the −9 and −8 positions based on the regression coefficients learned by the elastic net for PBSL=7 (Supplementary Fig. S4). When fewer than six G/C’s were found in the [-7,-1] range, PBSL between 11 and 15 had on average higher editing efficiency than other PBSL (Fig. 3D; highest median edit percentage of 9.8% when PBSL=13). Given a target site, these findings thus provided a general guideline for determining the optimal PBSL based solely on the 9 bp-long sequence upstream of the nick site in the edited-strand.

Some patterns of regression coefficients were shared among most combinations of PBSL and RTTL. Based on the magnitude of regression coefficients, the most salient nucleotides were those positioned around the nick site (Supplementary Fig. S4): C and G nucleotides at locations from −1 to +3, immediately 5’ to the PAM site, led to high editing efficiency, whereas T in the range from −2 to +3 were to be avoided. G/A at the −17 position was positively correlated with editing efficiency, while T at the same position was negatively correlated. In general, C at the +20 position was positively correlated with editing efficiency across designs. For the PAM site, AGG was favored and CGG disfavored for optimizing editing efficiency. Immediately after the PAM site, G at +7 position was strongly anti-correlated with editing efficiency. Surprisingly, the presence of G at +10 position was significant and tended to increase editing efficiency not only for RTTL=10, in which case G would be the last templated nucleotide as described above, but also for RTTL=15 or 20 when PBSL ≥ 11 (Fig. 3E). Considering that Cas9 variants similar to SpCas9 in size were known to recognize PAM sequences of varying length between 2-8 nts (57) and that the +10 position was only 4 bps downstream of the NGG PAM site, the fact that the presence of G at the +10 position consistently increased the editing efficiency across multiple RTTLs suggested that SpCas9 preferentially interacted with this specific nucleotide. Another unexpected finding was that select edited strand nucleotides even outside the regions of protospacer, PBS, and RT template seemed to modulate editing efficiency. For example, when PBSL=13 and RTTL=15, the presence of A at the +17 position, lying outside the range of the RT templated region, tended to decrease editing efficiency (Fig. 3F).

In summary, the elastic net regression models trained on each combination of PBSL and RTTL learned distinct and common sequence features modulating editing efficiency. The effect of the last templated G nucleotide on editing efficiency depended on both PBSL and RTTL of pegRNA designs. Moreover, pegRNAs with short PBSL highly depended on the GC content of PBS region, perhaps to help stabilize the hybridization to the complementary DNA. The PBS RNA-DNA hybridization energy was significantly anti-correlated with edit percentage in all combinations of PBSL and RTTL, but pegRNAs with shorter PBSL showed more pronounced dependence. Finally, the presence of G/C nucleotides was most critical immediately 5’ to the PAM site for all combinations of PBSL and RTTL.

### Deep neural network accurately predicts PE2 efficiency and yields interpretable features

It is challenging to model nonlinear effects of coupled nucleotides at multiple positions using elastic net regression. To extend our linear regression approach, we thus trained a deep neural network (DNN) model on Kim *et al.*’s data (25) to predict the edit percentage of a pegRNA design given its length and the sequence of the target site and flanking regions (Fig. 4A; Methods). The Pearson correlation between the observed and predicted edit percentages in the test set was 0.73 (Fig. 4B; Methods).

**Figure 4.**
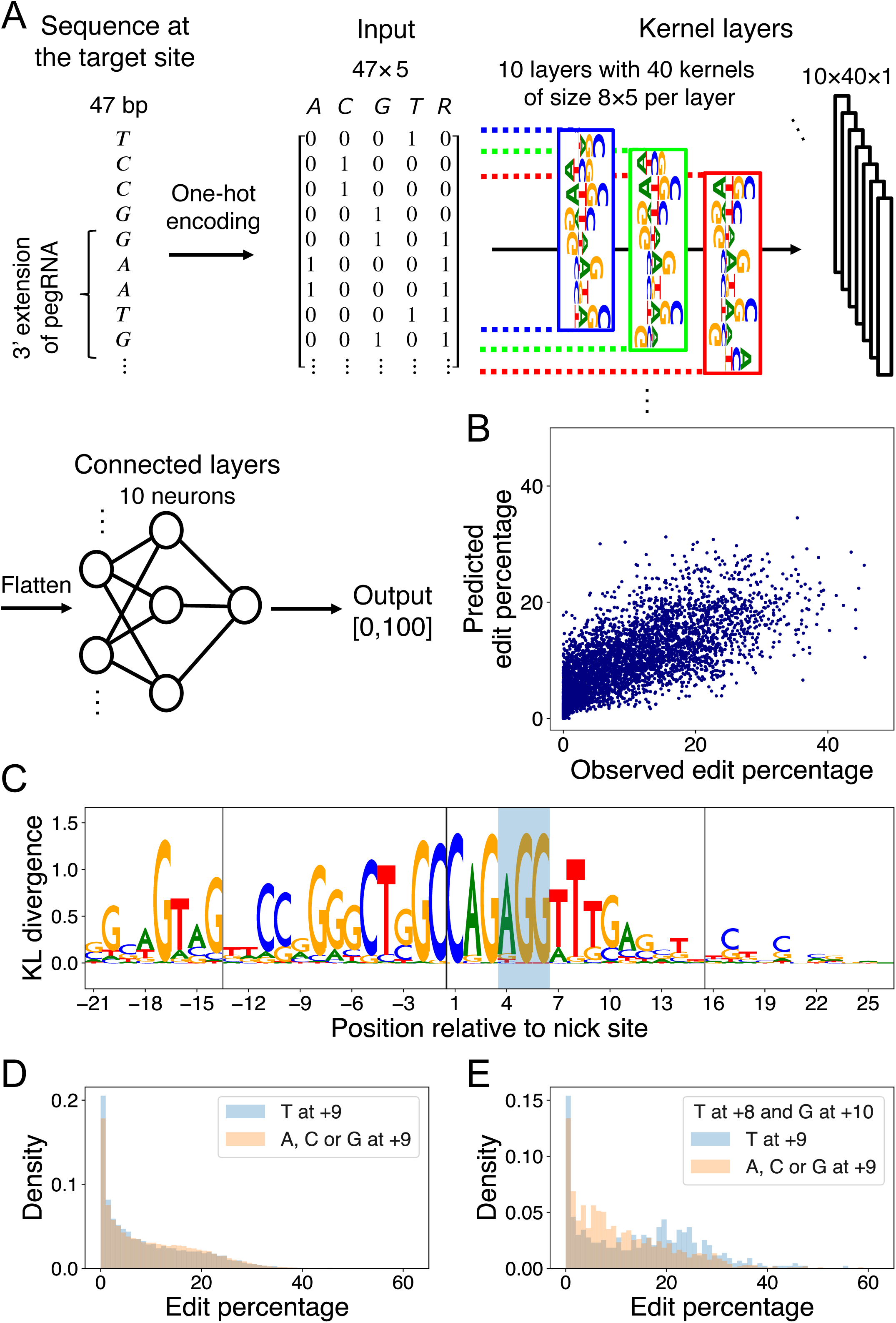
DNN learns marginal and multi-nucleotide sequence features. (A) DNN architecture. The first 4 columns of the input matrix indicate the presence of a particular nucleotide in the range [-21,+26] relative to the nick site via one-hot encoding; the 5^th^ column indicates whether a particular position resides in the union of the PBS and RT template regions of the pegRNA being considered. The input is passed through a layer of kernels and two dense layers to yield an output value between 0 and 100. (B) Scatter plot of observed versus predicted edit percentages of the target sites in the test set (Pearson *r* = 0.73, *p* < 10^-300^). (C) Position-wise KL divergence of maxEnt output nucleotide distributions with respect to the uniform nucleotide distribution (Supplementary Method S4). The vertical lines denote the borders of the PBS and RT template regions, and the PAM site is shaded. (D) Histograms of marginal edit percentages of the target sites containing T at +9 (blue) versus A, C, G at +9 (orange). (E) Same as in (D), but the target sites are conditioned to have T at +8 and G at +10.

To understand optimal sequences associated with high edit percentages, we applied a simulated annealing (SA) method for maximizing the prediction of a trained DNN over its input space via Markov Chain Monte Carlo (MCMC) sampling (41). We accelerated the simulation by using a broadened sampling distribution stemming from nonextensive statistical mechanics (42) and obtained the target sequences maximizing the predicted edit percentages for each combination of PBSL and RTTL (Supplementary Method S3; Supplementary Tables S11-S14). To identify the DNN-learned salient features in these optimal sequences, we next applied maxEnt, another MCMC sampling method based on the maximum entropy principle for generating new input sequences that produce similar DNN predictions as those of the initial input sequence (43). Upon initializing the maxEnt chains at the optimal target sequences, nucleotide preferences important for maximizing PE efficiency were extracted by calculating the position-wise Kullback-Leibler (KL) divergence between the nucleotide distribution in the sampled sequences and the uniform null distribution; in this formalism, the larger the KL divergence, the more relevant the nucleotide (Fig. 4C, Supplementary Fig. S7; Supplementary Method S4).

The DNN confirmed the sequence features previously detected by the elastic net model. For example, it learned that C at the +1 position, G at the +3 position, and G at the −17 position relative to the nick site were associated with high edit percentage; other similarities with the elastic net results included the preferences for A as the first nucleotide in the PAM sequence, for C or G in the PBS region, and for C at the +20 position. In addition, the DNN also discovered new features. For example, although elastic net detected no strong preference for any particular nucleotide at position +9, except for perhaps a weak preference for G (Supplementary Fig. S4), the DNN found T at this position to be associated with optimal editing. This discrepancy in the feature importance arose from the difference in model architectures; that is, the elastic net model learned only the independent effects of individual nucleotides, whereas the DNN accounted for aggregate effects of nucleotides captured by the filters. When considering only the marginal effect of single nucleotides at the +9 position, target sequences with T at that position actually had a lower edit percentage on average than those with other nucleotides at that same position (9.00% for T vs. 9.72% for A,C,G at +9; Fig. 4D). However, imposing the +8 position to be T and the +10 position to be G showed that target sequences with T at +9 had a higher edit percentage on average than those with other nucleotides at +9 (14.33% for T vs. 10.72% for A,C,G at +9; Fig. 4E; Supplementary Fig. S8A,B). Similarly, the marginal effect of T at the −4 position, residing in the PBS region, was negligible; however, the DNN uncovered a role of T in the context of GC-rich background: for sequences with 100% GC content in the range [-7,-1] except at the −4 position, the presence of T at −4 substantially increased the edit percentage by 6.90% on average (Supplementary Fig. S8C,D). Finally, the DNN found a preference for CAG at [+1,+3] in optimal target sequences, whereas elastic net found only a marginal effect of G/C at +2 and a less pronounced trinucleotide effect of C(G/C)G in the same region (Supplementary Fig. S8E,F). In summary, our DNN was able to predict the observed edit percentage with high accuracy and yielded interpretable results regarding both marginal and context-dependent sequence features of optimal target sites.

## DISCUSSION

We have demonstrated that both regional heterochromatin and local nucleosome occlusion of target sites may decrease PE2 editing efficiency, with a more pronounced effect observed for the H3K9me3 modification, perhaps because unwinding DNA may be particularly difficult in a heterochromatin environment where multiple nucleosomes are condensed together. These results are consistent with a recent report showing that inducing open chromatin state improves PE and base editor efficiencies (58). Our study thus provides evidence for the hypothesis that chromatin structure modulates genome editing efficiency by interfering with the accessibility of target sites to Cas9 protein and guide RNA. Unlike the significant predictive power of H3K9me3 in classifying strongly and weakly editable target sites, however, the feature of H3K27me3 was insignificant (Supplementary Fig. S2A,B; Supplementary Table S3). While both H3K9me3 and H3K27me3 are associated with heterochromatin formation, H3K9me3 is associated with constitutive heterochromatin, whereas H3K27me3 is associated with facultative heterochromatin (59) which depends on the presence of certain stimuli and thus remains conducive to dynamic changes (60). Further investigation is needed to decipher the precise differences in folding pattern making constitutive heterochromatin more resistant to editing than facultative heterochromatin.

Our work shows that, in addition to the chromatin environment, local sequence content of target sites can also modulate editing efficiency. For example, strategic positioning of G and C nucleotides in the PBS and RT regions increases the editing efficiency, indicating that G:C base pairing between DNA and pegRNA helps anchor the pegRNA to the target site prior to editing. A similar observation regarding the GC content in the PBS region has been previously reported (25); however, our study further clarifies that 1) the presence of G or C is most important in the RTT region, immediately downstream of the PBS region, at positions +1 and +3, 2) within the PBS region itself, the positive effect of G or C is most pronounced near the nick site and rapidly decays away from the nick site for PBSL ≥ 11 (Fig. 3A, Supplementary Fig. S4), and 3) the importance of GC content depends on PBSL (Supplementary Fig. S5), with the effect size being particularly strong for pegRNAs with shorter PBSL, which may need more G:C base pairs to compensate for weak overall PBS-DNA hybridization interactions of only a small number of bases. As previously observed (14), we find that having G as the last templated nucleotide may decrease editing efficiency; but, we show that this effect holds only in pegRNAs with sufficiently long PBSL and RTTL. Kim *et al*. have similarly observed that for RTTL≤12, editing efficiency is on average the highest if the last templated nucleotide is G (25); but, we show that this positive effect may be unrelated to G being the last templated nucleotide, as G at the +10 position has a positive correlation with editing efficiency regardless of the RTTL. Considering that the +10 position is only 4bp downstream of the canonical NGG PAM site, it is plausible that the G nucleotide at this location affects the binding of Cas9.

When choosing a target site, our results recommend selecting target sequences that (i) have AGG in the PAM site, (ii) have high GC content in the PBS region especially when the PBS is short, (iii) avoid G as the last templated nucleotide if the RT template is longer than 12, and (iv) have G at the −17 position relative to the nick site on the edited strand (Fig. 5). Once the target site has been chosen, a PBSL in the [11,13] nt range is generally recommended, unless the target site has high GC content that provides stable hybridization between the pegRNA and target DNA, in which case a shorter PBSL is recommended. As for the effect of RTTL, pegRNAs with shorter RTTL have on average higher editing efficiency than those with longer RTTL; when the available PAM site is far away from the desired site, such that having RTTL > 15 nt is unavoidable, we recommend the following designs: prioritize the last templated nucleotide in the order T≈C>A>G. Short pegRNAs are generally advantageous, as long as the PBS region has sufficiently high GC content to hybridize stably with its target DNA (Fig. 5).

**Figure 5.**
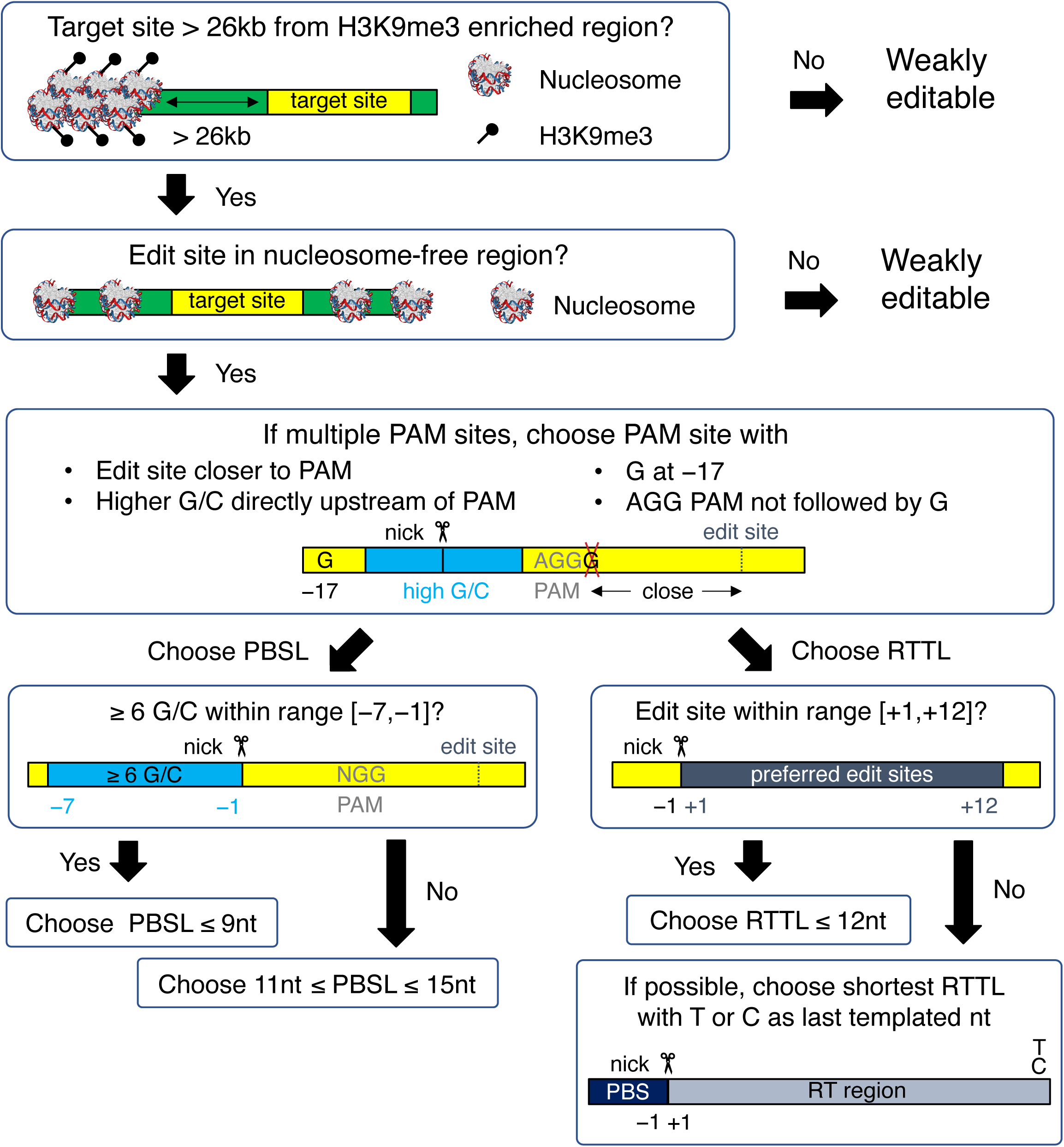
Flow chart for designing an optimal pegRNA for selected target site. The yellow boxes represent the target site, which is defined to be the union of edited-strand nucleotides spanning the protospacer region, the PBS region, and the RT region. The green boxes represent DNA flanking the target site. All nucleotide numbers are relative to the nick site.

Our computational analysis has identified several epigenetic and sequence features that need to be considered when designing pegRNAs to optimize PE efficiency. Future availability of additional high-throughput genome editing data and biophysical studies investigating how PEs search and bind target sequences will help further improve our understanding and make this technology feasible for effective biomedical applications.

## Supporting information

Supplementary Methods, Figures, Tables, and References

## DATA AVAILABILITY

Source code used to generate the results is available at https://github.com/jssong-lab/HOPE.

## ACKNOWLEDGMENT

Part of this project was completed while J.S.S. was on sabbatical leave in the Center for Theoretical Physics at the Massachusetts Institute of Technology and the Department of Statistics at Harvard University.

