## Supplementary Methods, Figures, Tables, and References for "Chromatin structure and context-dependent sequence features control prime editing efficiency"

**This PDF file includes:**

Supplementary Methods S1-S4
Supplementary Figures S1-S8
Supplementary Tables S1-S14
References for Supplementary Materials

### SUPPLEMENTARY METHODS

#### Supplementary Method S1: Editing efficiency datasets used in this study

PE2 target sites and the corresponding editing efficiencies were obtained from Kim *et al.* (1), Anzalone *et al.* (2), and in-house validation experiments. All editing experiments were performed in HEK293T cells. The dataset from Kim *et al.* included two subsets: 1) a set of experiments using 43149 different pegRNA designs targeting 1956 unique exogenous sequences integrated via a lentiviral plasmid library and containing variations in the PBS length (PBSL) and RT template length (RTTL) for each spacer sequence (Supplementary Table 4 from Kim *et al.*), and 2) a set of experiments using 32 pegRNA designs targeting 32 distinct endogenous loci in the human genome, without varying the PBSL and RTTL for each spacer sequence (Supplementary Table 3 from Kim *et al.*). The datasets from Anzalone *et al*. consisted of 88 different pegRNA designs targeting 8 endogenous loci with varying RTTL for each spacer sequence (Fig. 2b and Extended Data Fig. 5a-c from Anzalone *et al.*). The in-house validation experiments consisted of 25 pegRNA designs targeting 25 distinct endogenous loci, without varying the PBSL and RTTL for each spacer sequence. Binary classification analysis for determining the effects of local and global chromatin structure was performed using only the endogenous target sites from Kim *et al.*, Anzalone *et al*. and in-house validation experiments. When replicates were available, editing efficiencies were determined by averaging the percentage of successful edits across the replicates (Methods). Regression analyses for determining the salient sequence features were performed using the integrated target sites from Kim *et al*. and endogenous sites from Anzalone *et al.*; the OLS linear regression model used both datasets, and the elastic net and deep neural network regression models used only Kim *et al.*'s dataset.

**Supplementary Method S2: Singular Value Decomposition (SVD) determines a consensus set of locations where multiple immuno-precipitated (IP) samples of H3K9me3 are enriched relative to input**

From the collection of read densities for 3 IP and 3 input samples partitioned into $L$ bins of size 1kb, we constructed a matrix $X$ of size $6 \times L$, where the rows and columns of $X$ represent the samples and bins, respectively. Matrix $X$ was decomposed as

$$X = USW^T = VW^T$$

where $U$ is the matrix of left singular vectors as columns decomposing the 6-dimensional sample space, $S$ is the diagonal matrix of singular values in descending order, and $W$ is the matrix of right singular vectors as columns decomposing the row space $span(\{X_{i:}|1 \leq i \leq 6\})$, a 6-dimensional subspace of the $L$-dimensional location space. This SVD decomposition represented the $i$th sample, $X_{i:}$, as a 6-dimensional vector $V_{i:}$ using $\{W_{:j}|1 \leq j \leq 6\}$ as a basis of the row space. We then determined the locations of differential enrichment between IP and input samples by comparing $\{V_{i:}|i \in \mathrm{IP}\}$ and $\{V_{i:}|i \in \mathrm{input}\}$ (Supplementary Fig. S1A). We calculated the variance ratio $v_j$ of the two groups $\{V_{ij}|i \in IP\}$ and $\{V_{ij}|i \in \mathrm{input}\}$ as the ratio of between-group variance to within-group variance. The variance ratios $v_1$, $v_2$, and $v_3$ had the highest values among the 6 components of $v$ (Supplementary Fig. S1B), and we thus defined an $L$-dimensional location vector $y$ as a weighted linear combination of $W_{:1}, W_{:2},$ and $W_{:3}$, such that $y$ reflected the enrichment of the read densities of the IP group relative to those of the input group. The coefficients $\alpha_j$ of the linear combination of $W_{:1}, W_{:2},$ and $W_{:3}$ were computed via the equation

$$\alpha_j = \frac{1}{N_{\mathrm{IP}}} \sum_{i \in \mathrm{IP}} V_{ij} - \frac{1}{N_{\mathrm{input}}} \sum_{i \in \mathrm{input}} V_{ij} \quad \text{for } j = 1,2,3,$$

where $N_{\mathrm{IP}}$ and $N_{\mathrm{input}}$ are the the number of samples in the IP group and input group, respectively.

Adjacent bins corresponding to the top 20% of positive components of $y$ were aggregated. All aggregated bins that contain at least one bin from the top 5% of positive components of $y$ were determined to be the consensus regions of enrichment between H3K9me3 IP read densities and input read densities (Supplementary Fig. S1C).

**Supplementary Method S3: Simulated annealing using the Tsallis-Boltzmann distribution discovers sequences that maximize deep neural network (DNN) predictions of PE efficiency**

The simulated annealing (SA) algorithm is a time-inhomogeneous Markov Chain Monte Carlo (MCMC) sampling method designed to find the global extremum of a function without getting trapped in local extrema (4). Tsallis *et al.* proposed an accelerated version of SA based on a sampling distribution with a broadened tail, arising from the nonextensive statistical mechanics formalism of maximizing the generalized *q*-entropy (5). We used this modified SA algorithm to maximize $EP$, the edit percentage predicted by our DNN, as a function of the pegRNA target sequence and its flanking sequences. At each iteration step $i$ in the Markov chain, the probability $p$ of accepting a symmetric proposal transition from sequence $s_{\text{previous}}$ to sequence $s_{\text{proposed}}$ was calculated in terms of the Tsallis-Boltzmann distribution as

$$p(s_{\text{previous}} \rightarrow s_{\text{proposed}}) = \begin{cases} 1, & \Delta EP \geq 0 \\ \left(1 - (q_A - 1)\frac{\Delta EP}{T_i}\right)^{-1/(q_A-1)}, & \Delta EP < 0 \end{cases}$$

where $\Delta EP = EP(s_{\text{proposed}}) - EP(s_{\text{previous}})$ and $q_A$ is a chosen parameter constrained to values $> 1$. The variable $T_i$, a term analogous to the temperature parameter in classical SA, was scheduled to decrease at every iteration step according to the equation

$$T_i = T_1 \frac{2^{(q_V-1)}-1}{(1+i)^{(q_V-1)}-1},$$

where $T_1$ and $q_V$ are chosen parameters constrained to values $> 0$. The acceptance probability $p$ and the cooling schedule for $T_i$ converge to their classical SA form as $q_A$ and $q_V$ approach the value $1$. The advantage of using the above acceptance probability and cooling schedule over those in classical SA is that for larger values of $q_A$ and $q_V$, respectively, the acceptance distribution becomes broader and the cooling rate more rapid, enabling the modified SA to explore a wider area in the sequence space and converge faster to the global maxima.

For each Markov chain, the GG dinucleotide in the PAM motif and the indicator variables encoding for the locations of the PBSL and the RTTL in the 47 nt-long sequence were fixed. In each iteration step of the modified SA, one uniformly chosen base from the remaining 45 nts in $s_{\text{previous}}$ was proposed to be mutated into a new nucleotide with 1/3 probability for the three possible substitutions. For each combination of PBSL and RTTL, we chose 25 sequences from the test set which produced the highest predicted edit percentages and initialized the SA algorithm using each of these sequences; from the resulting 25 approximate maxima found by SA, we chose the sequence yielding the highest predicted edit percentage for further analysis, as described in Supplementary Method S4.

To determine an appropriate value of $T_1$, for each combination of PBSL and RTTL, we initialized $5 \times 10^5$ random 47 nt-long DNA sequences with a GG dinucleotide fixed at the 25$^{\text{th}}$ and 26$^{\text{th}}$ position. For each initialized sequence, we mutated a single uniformly chosen base as described above and computed the absolute value of the difference in $EP$ between the mutated and initialized sequences. We set $T_1 = 5$, which was the integer closest to the 99$^{\text{th}}$ percentile of absolute values of changes in $EP$. To determine an appropriate choice of $q_A$ and $q_V$, we restricted ourselves to the case when PBSL=13 and RTTL=15 and performed a grid search for $q_A$ and $q_V$ both in the range [1,1.6], stepping coarsely at first with step size 0.2 and then more finely with step size 0.05 in the region where SA was preforming better (Supplementary Tables S11-S14). For each pair of $q_A$ and $q_V$, we simulated 10 independent Markov chains with $2 \times 10^6$ iterations, where the chains were initialized with the same 10 random sequences for all pairs of $q_A$ and $q_V$. For each chain, we measured $i_{max}$, the number of iteration steps required to achieve the first observation of $EP_{max}$, the maximum $EP$ found within that chain. We then calculated the average $EP_{max}$ and $i_{max}$ across the 10 chains for each pair of $q_A$ and $q_V$. We chose $q_A = 1.15$ and $q_V =$

$1.30$, because the modified SA using this combination consistently required the fewest iterations compared to other combinations of $q_A$ and $q_V$ to find optimal sequences with $EP$ greater than or equal to the $EP_{max}$ of classical SA (Supplementary Tables S11-S14) .

**Supplementary Method S4: Identifying salient features of optimal pegRNA sequences**

To determine the important nucleotides and positions in the sequences predicted by our DNN to have high editing efficiency, we applied maxEnt, a statistical interpretation method using MCMC sampling of new input sequences from the Boltzmann distribution associated with a trained DNN. In this formalism, the energy of each sample is given by its distance from a fixed initial sequence, measured with respect to the $\ell_2$-norm on the penultimate activation space of DNN; by design, the MCMC samples preserve salient features important for DNN prediction, while replacing unimportant positions with random nucleotides (6). For each pair of PBSL and RTTL, maxEnt was initialized using the sequence found by SA to have the highest predicted edit percentage. In each iteration step of MCMC sampling, one uniformly chosen base from the 47nt-long sequence, excluding the GG dinucleotide in the PAM motif, was proposed to be mutated into a new nucleotide, with 1/3 probability for the three possible substitutions. The temperature was chosen such that the acceptance rate of the first iteration step was 0.10. Each initial sequence was seeded into 1000 independent Markov chains with $10^5$ iterations, and the simulation results were analyzed as follows: After discarding the first 10% of iterations as burn-in, sequences subsampled at every $10^3$ iterations were aggregated, and their one-hot-encoded matrices were averaged across all 1000 chains, resulting in a position-specific score matrix (PSSM). Salient sequence features were determined by computing the position-wise Kullback-Leibler (KL) divergence of the resulting PSSM with respect to the uniform distribution.

### SUPPLEMENTARY FIGURES

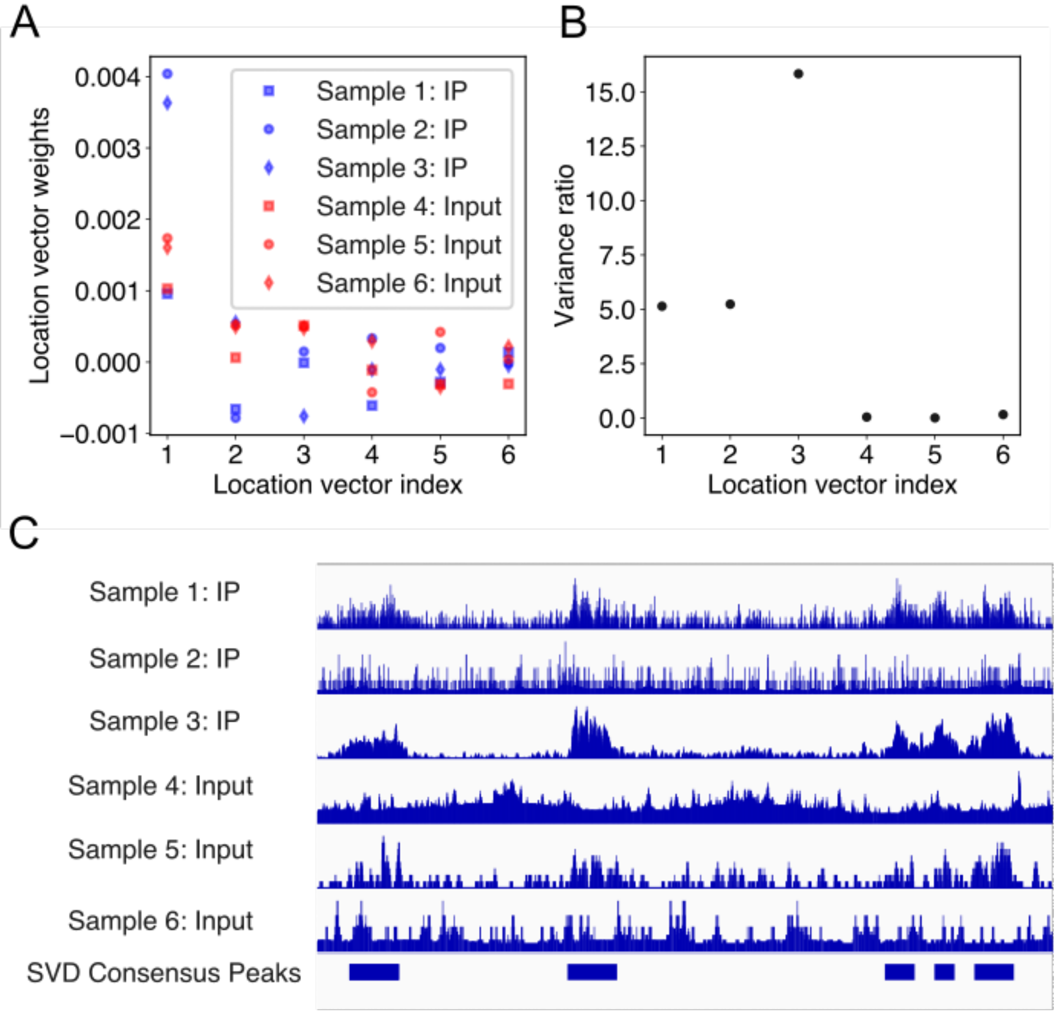


**Figure S1:** Integration of multi-sample histone modification data. The whole genome was partitioned into $L$ bins, and the H3K9me3 ChIP-seq read density for the $i$th sample was expressed as an $L$-dimensional row vector $X_{i:}$. The data vectors $X_{i:}$ were then simultaneously decomposed as $X = VW^T$ (Supplementary Method S2). (A) Matrix elements $V_{ij}$, which weights the $j$th location vector for the $i$th sample. Blue points represent the IP samples, and red points represent the input samples. The different shapes of points represent different samples. (B) Variance ratio of all vector elements of $V_{:j}$ between IP and input groups. (C) Example of consensus peaks determined by taking a linear combination of the first three location vectors (Methods, Supplementary Method S2).

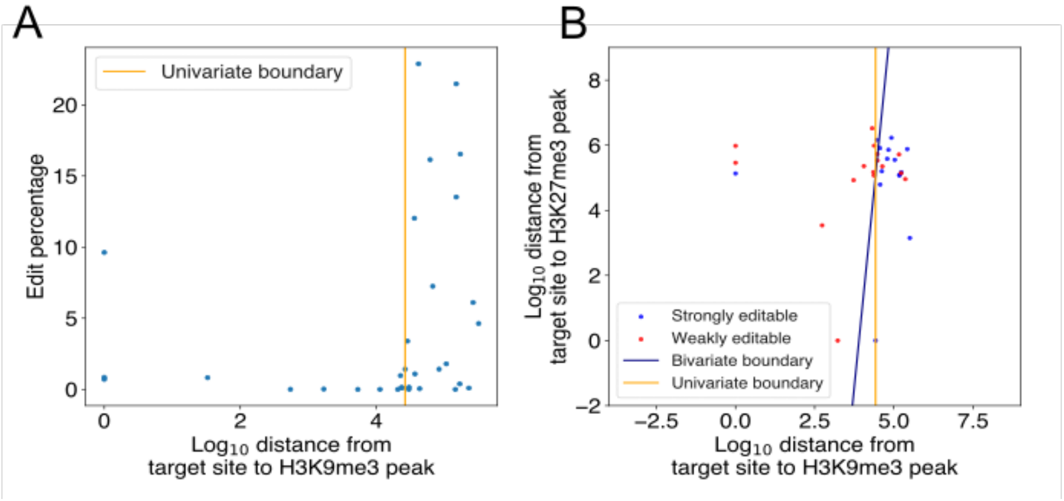


**Figure S2:** Classification of edit percentages based on epigenomic characterizations of target sites. (A) Scatter plot of edit percentage versus $\log_{10}$ distance of target site to the nearest H3K9me3 peak in the training set consisting of 32 endogenous target sites from Kim *et al.* (Methods, Supplementary Method S1). The orange line refers to the logistic regression decision boundary. (B) Scatter plot of $\log_{10}$ distance of target site to the nearest H3K9me3 peak versus $\log_{10}$ distance of target site to the nearest H3K27me3 peak. Blue and red dots represent target sites strongly and weakly editable, respectively. The threshold for determining strongly and weakly editable target sites was 1%. The blue line represents the bivariate logistic regression decision boundary using both H3K9me3 and H3K27me3 features. The orange line represents the univariate logistic regression decision boundary using only H3K9me3.

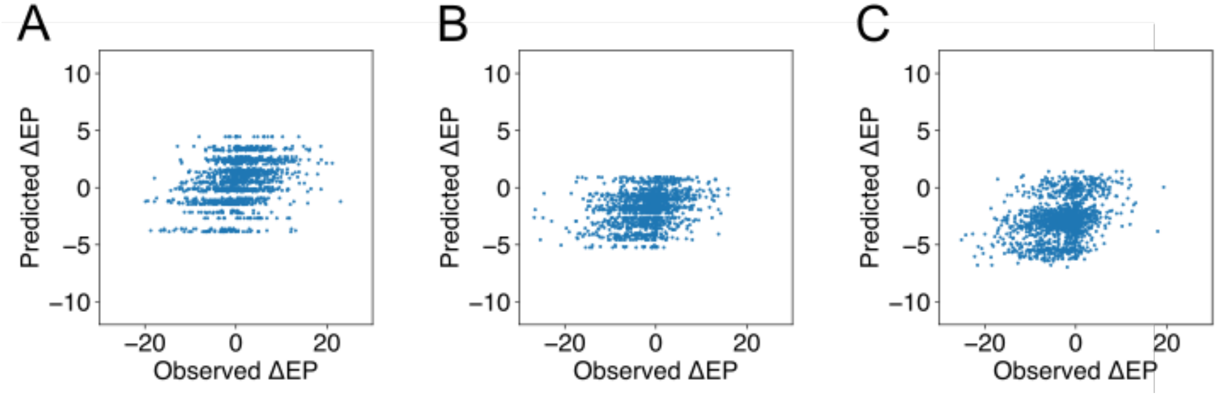


**Figure S3:** Performance of the OLS linear regression model on Kim *et al.*'s data derived from integrated target sites for RTTL in the range (A) [10,12] (Pearson $r = 0.34$, $p = 6.65 \times 10^{-41}$), (B) [12,15] (Pearson $r = 0.26$, $p = 2.86 \times 10^{-24}$), and (C) [15,20] (Pearson $r = 0.33$, $p = 1.20 \times 10^{-39}$).

A

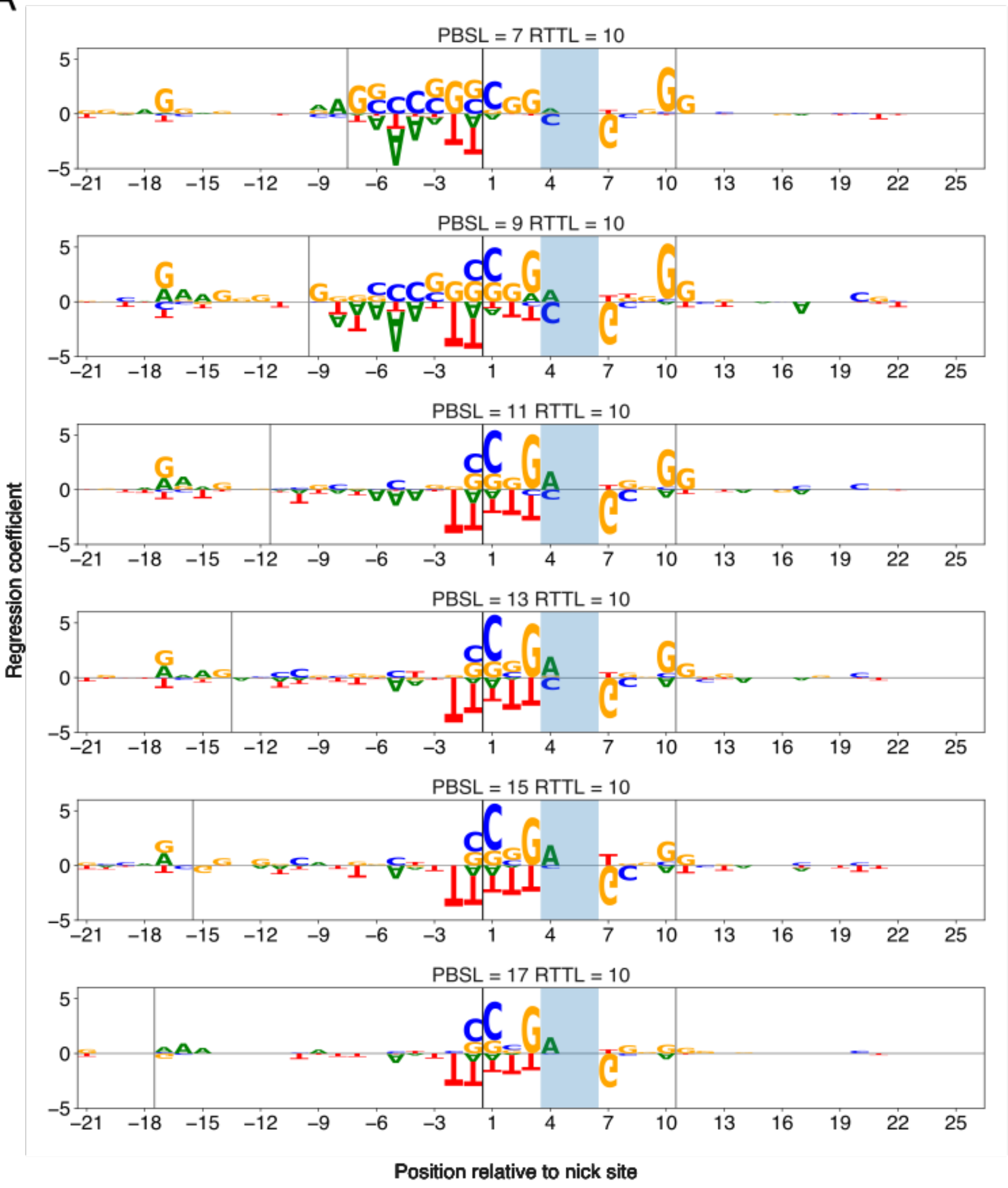
PBSL = 7 RTTL = 10
PBSL = 9 RTTL = 10
PBSL = 11 RTTL = 10
PBSL = 13 RTTL = 10
PBSL = 15 RTTL = 10
PBSL = 17 RTTL = 10
Regression coefficient
Position relative to nick site

B

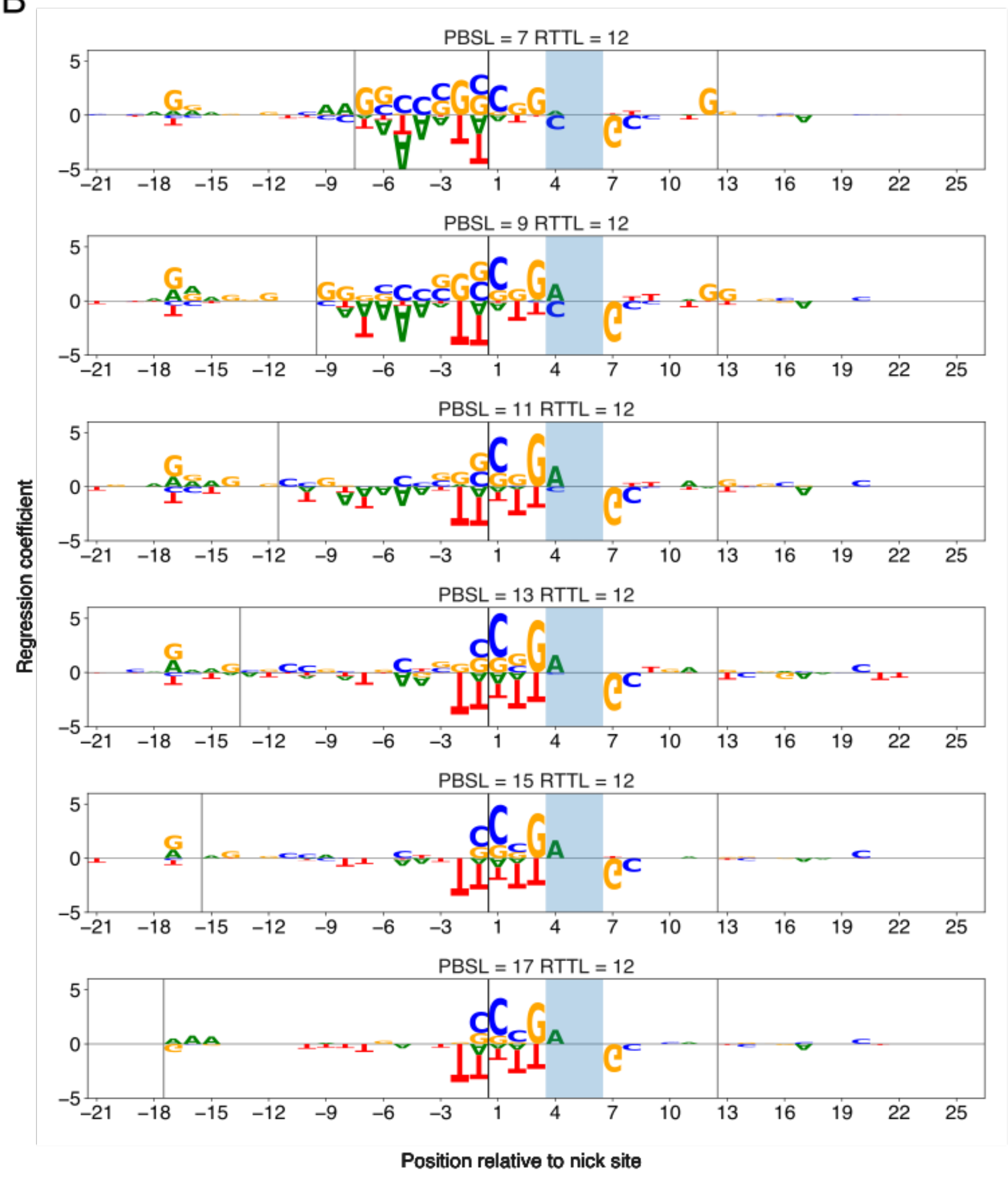

PBSL = 7 RTTL = 12
PBSL = 9 RTTL = 12
PBSL = 11 RTTL = 12
PBSL = 13 RTTL = 12
PBSL = 15 RTTL = 12
PBSL = 17 RTTL = 12
Regression coefficient
Position relative to nick site

C

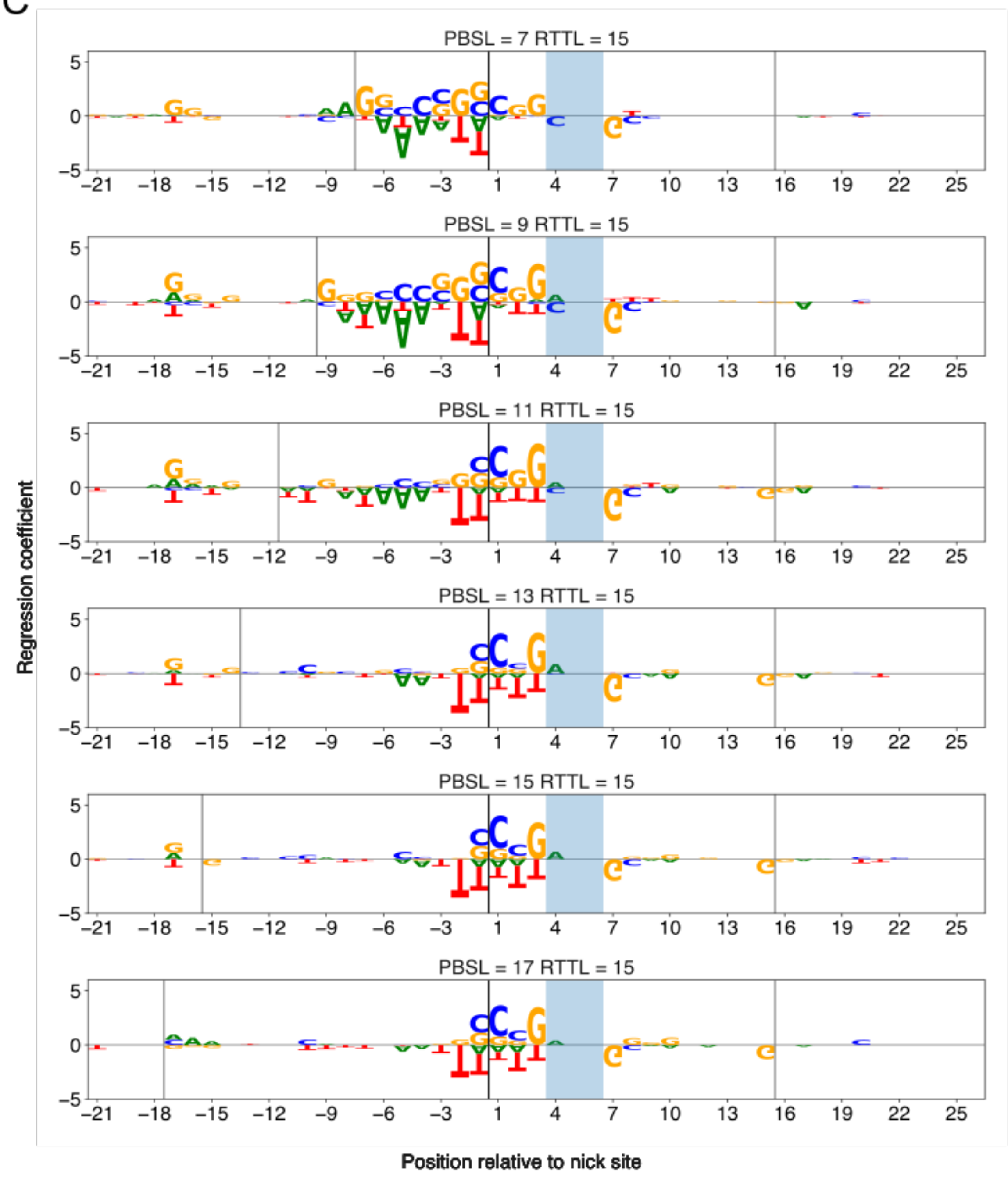
PBSL = 7 RTTL = 15
PBSL = 9 RTTL = 15
PBSL = 11 RTTL = 15
PBSL = 13 RTTL = 15
PBSL = 15 RTTL = 15
PBSL = 17 RTTL = 15
Regression coefficient
Position relative to nick site

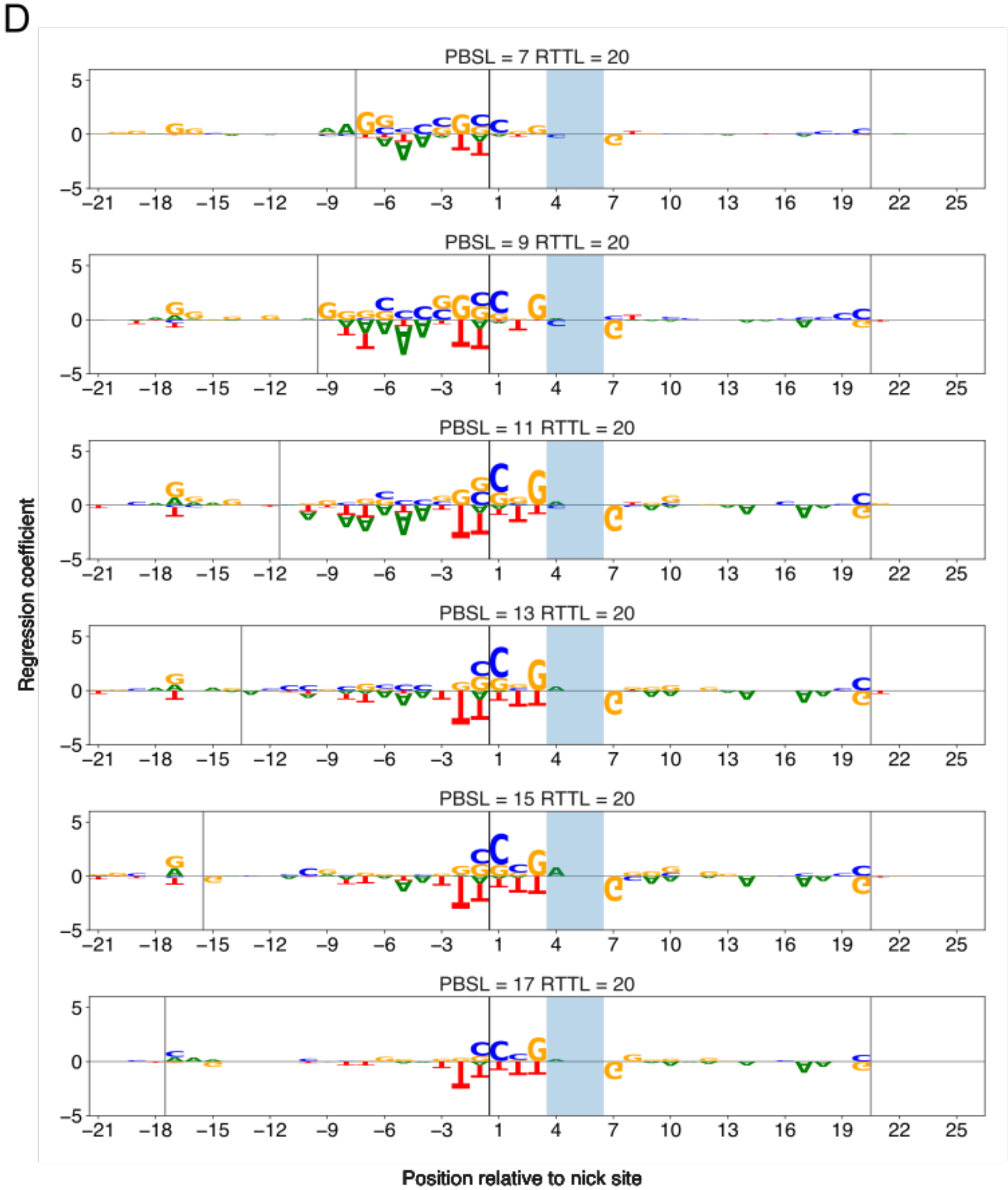


**Figure S4:** Representation of elastic net regression coefficients at positions in the range [-21,+26] with respect to the nick site when (A) RTTL=10, (B) RTTL=12, (C) RTTL=15, and (D) RTTL=20. The elastic net regression model was trained on Kim *et al.*'s data derived from integrated target sites. The letters indicate the nucleotides in the edited strand. Heights of the letters represent the absolute value of the regression coefficients. The vertical lines represent the delineation of PBS and RT regions, and the shaded region represents the PAM motif.

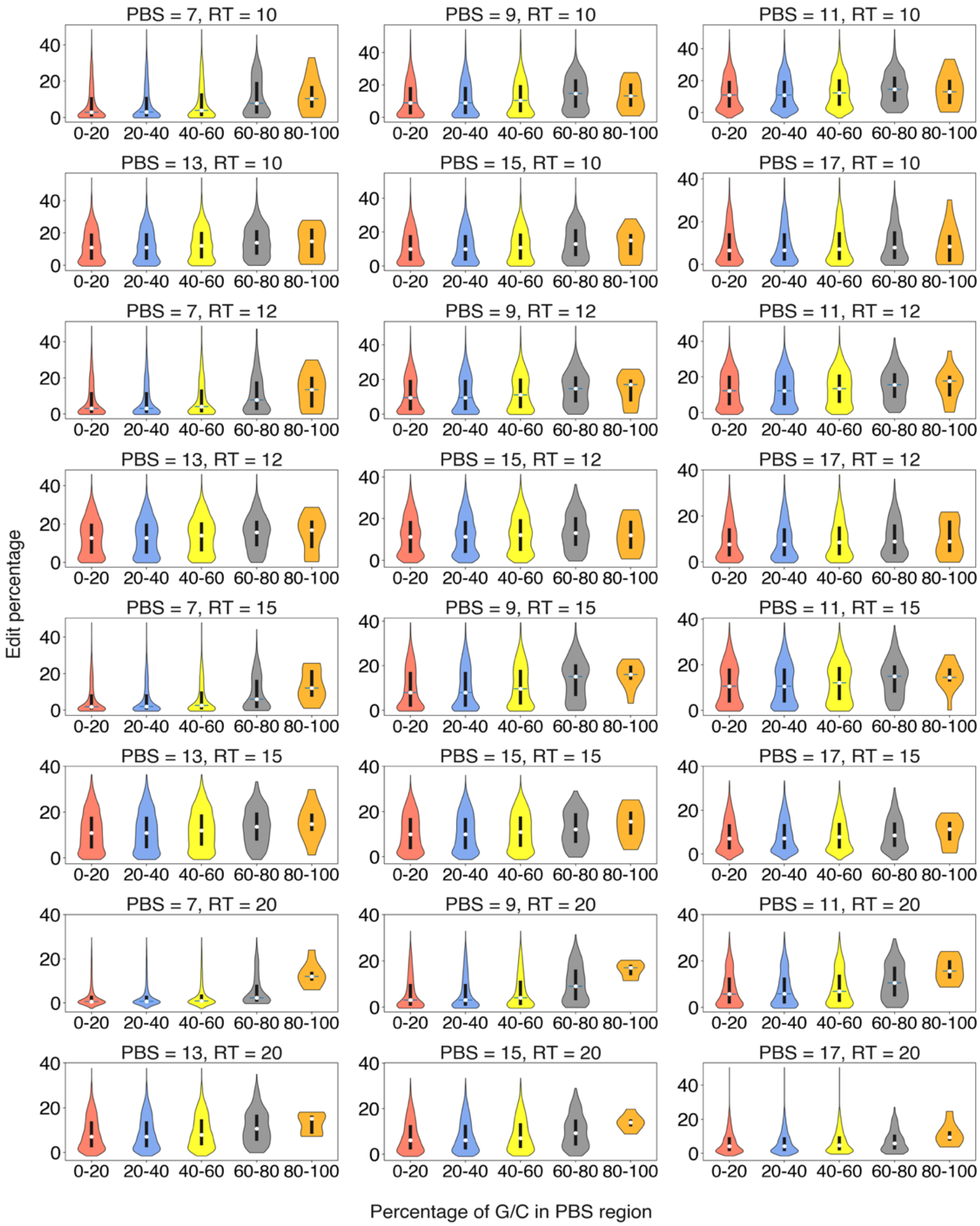


**Figure S5:** Violin plots of edit percentages for target sequences binned according to GC content in the PBS region. Plots were generated for all combinations of PBSL and RTTL available in Kim *et al.'s* data derived from integrated target sites. The white dots represent the median edit percentage, and the thick black lines represent the portion of the data between the first and third edit percentage quartiles.

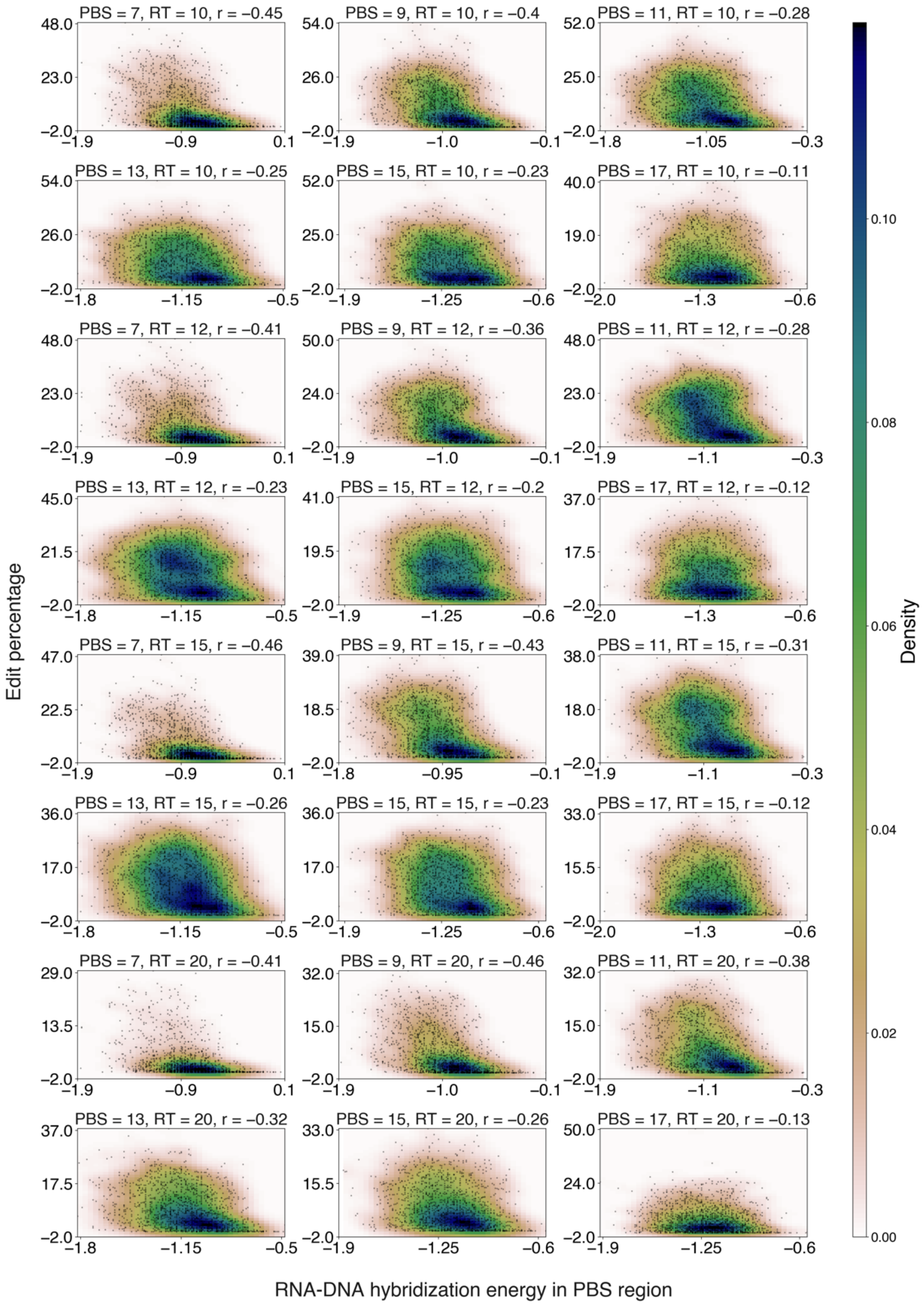


**Figure S6:** Scatter plots of edit percentage versus PBS-DNA hybridization energy for all combinations of PBSL and RTTL available in Kim *et al.*'s data derived from integrated target sites. The colors represent the density of data points in the scatter plot. Pearson $r$ is shown above each figure.

A

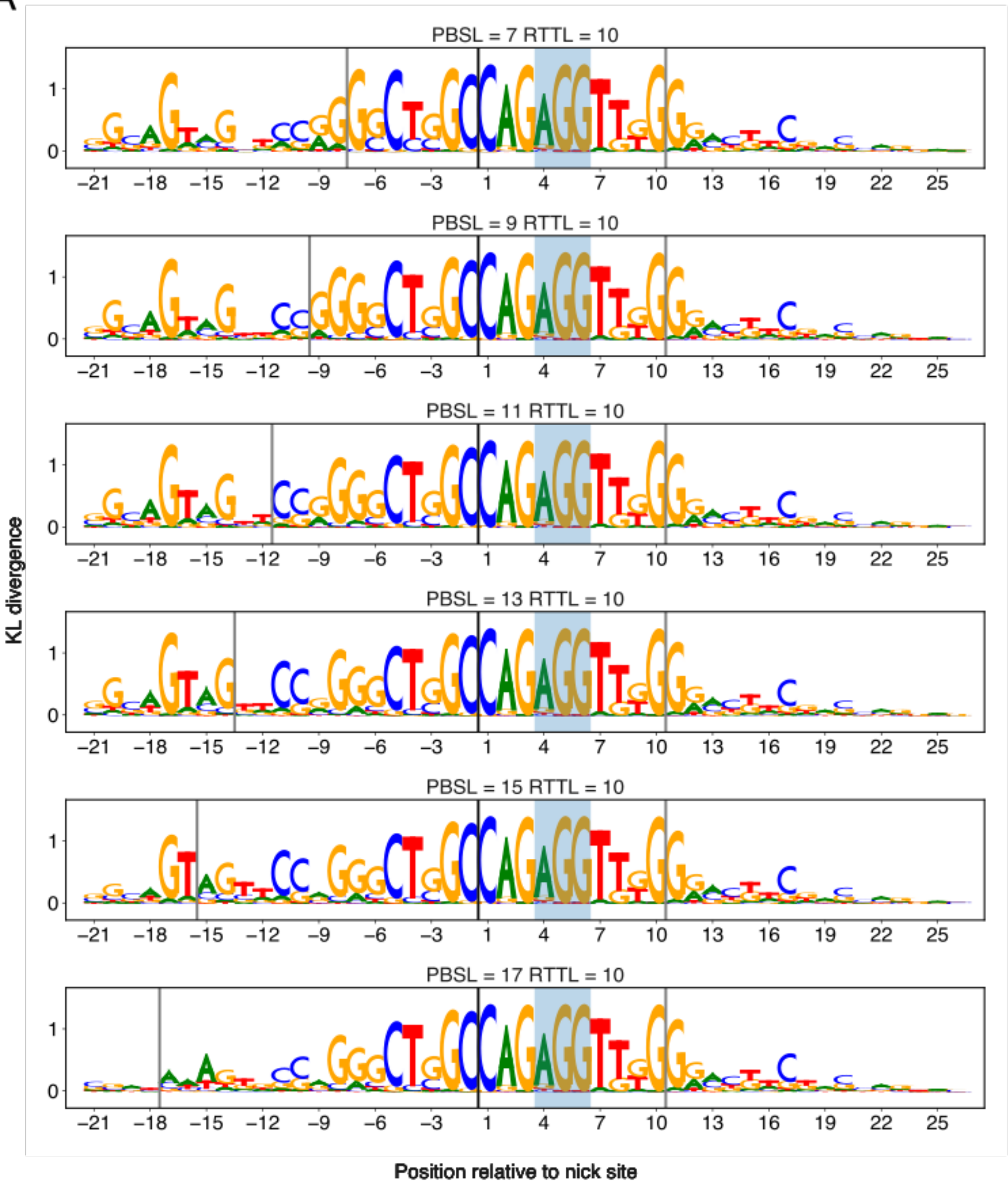
PBSL = 7 RTTL = 10
PBSL = 9 RTTL = 10
PBSL = 11 RTTL = 10
PBSL = 13 RTTL = 10
PBSL = 15 RTTL = 10
PBSL = 17 RTTL = 10
KL divergence
0
1
−21 −18 −15 −12 −9 −6 −3 1 4 7 10 13 16 19 22 25
Position relative to nick site

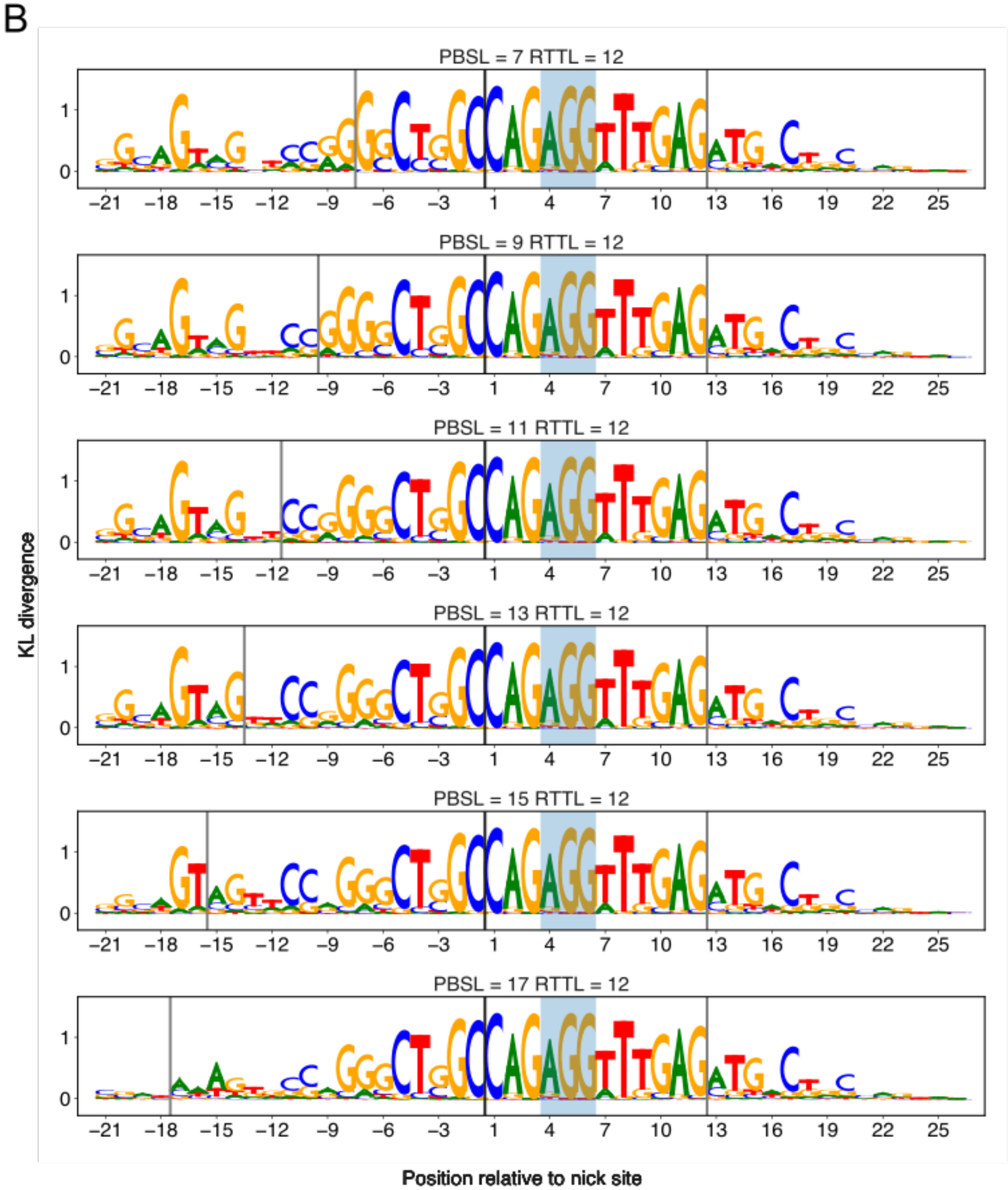

B
PBSL = 7 RTTL = 12
PBSL = 9 RTTL = 12
PBSL = 11 RTTL = 12
PBSL = 13 RTTL = 12
PBSL = 15 RTTL = 12
PBSL = 17 RTTL = 12
KL divergence
Position relative to nick site

C

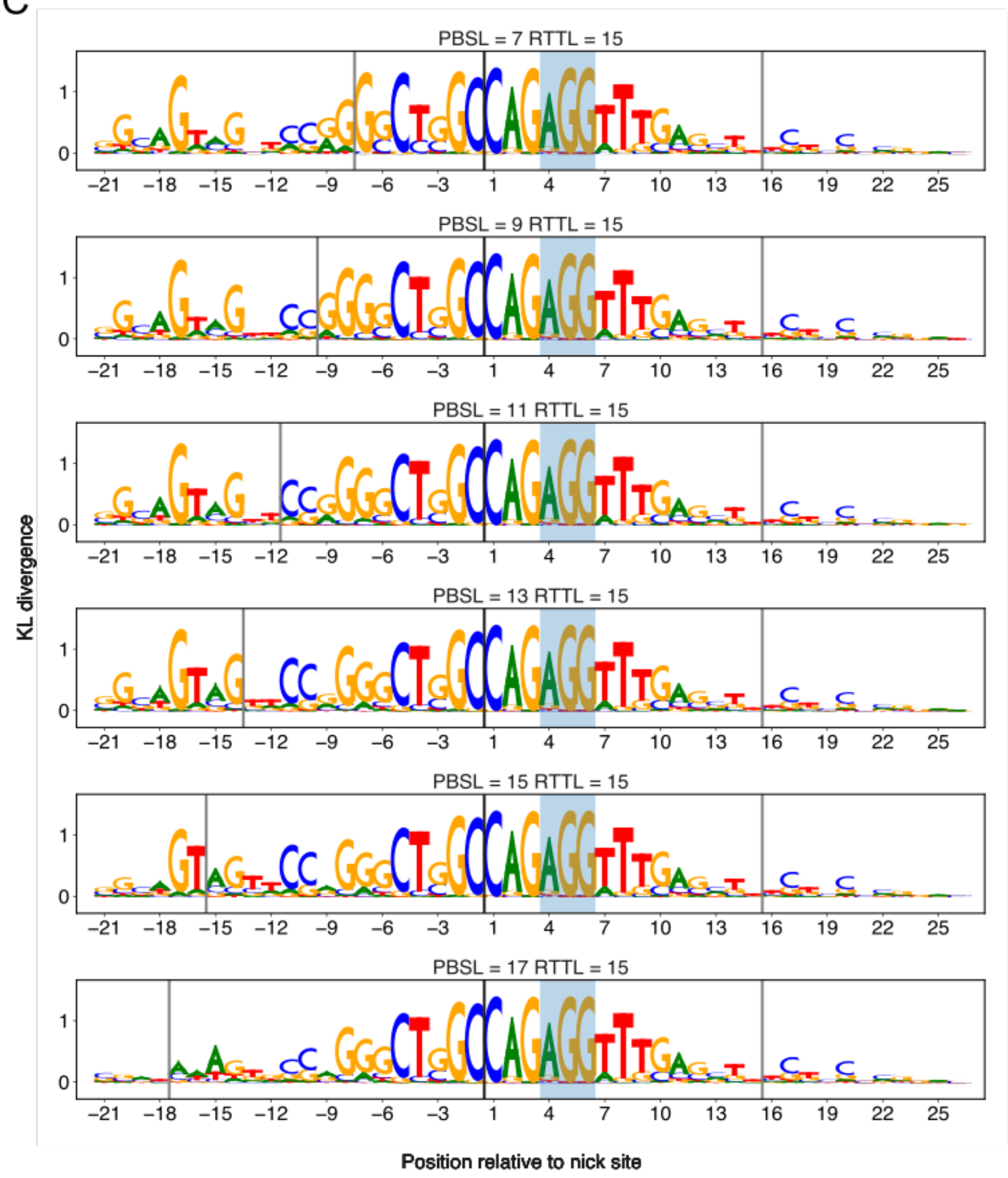

PBSL = 7 RTTL = 15
PBSL = 9 RTTL = 15
PBSL = 11 RTTL = 15
PBSL = 13 RTTL = 15
PBSL = 15 RTTL = 15
PBSL = 17 RTTL = 15
KL divergence
Position relative to nick site

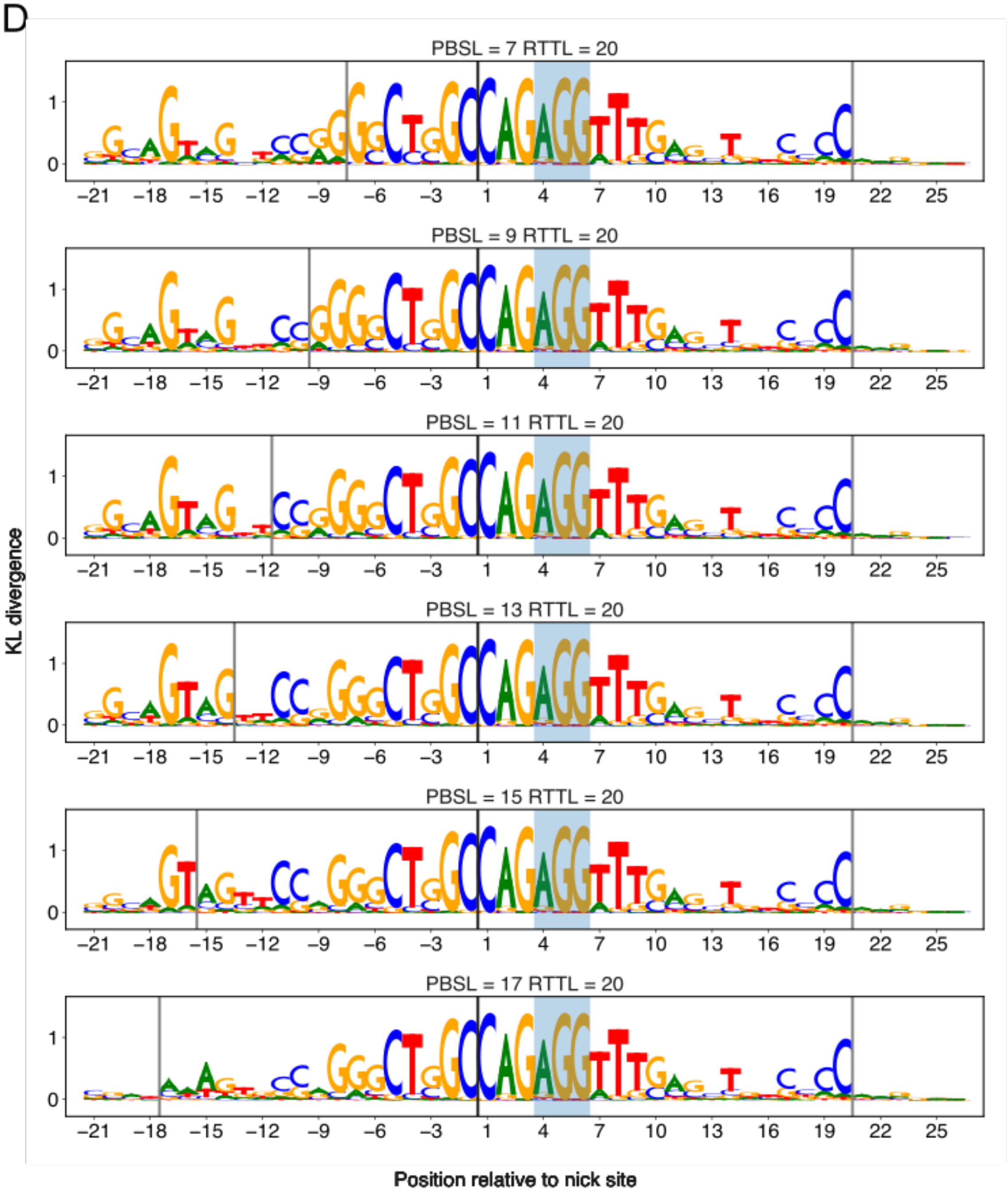


**Figure S7:** Sequence logos of maxEnt samples. Position-wise KL divergence of maxEnt outputs with respect to the uniform distribution are shown for (A) RTTL=10, (B) RTTL=12, (C) RTTL=15, and (D) RTTL=20. Positions of the KL divergence are labeled starting from the -21 position to the +26 position relative to the nick site. The vertical lines represent the delineation of PBS and RT regions, and the shaded region represents the PAM motif. The maxEnt outputs were initiated from the sequences found by simulated annealing (Supplementary Method S3) to maximize the deep neural network trained on Kim *et al.'*s data derived from integrated target sites (Methods; Supplementary Method S4). The letters indicate the DNA nucleotides in the edited strand, and the heights of the letters at each position are proportional to the relative nucleotide frequencies at that position in the maxEnt samples.

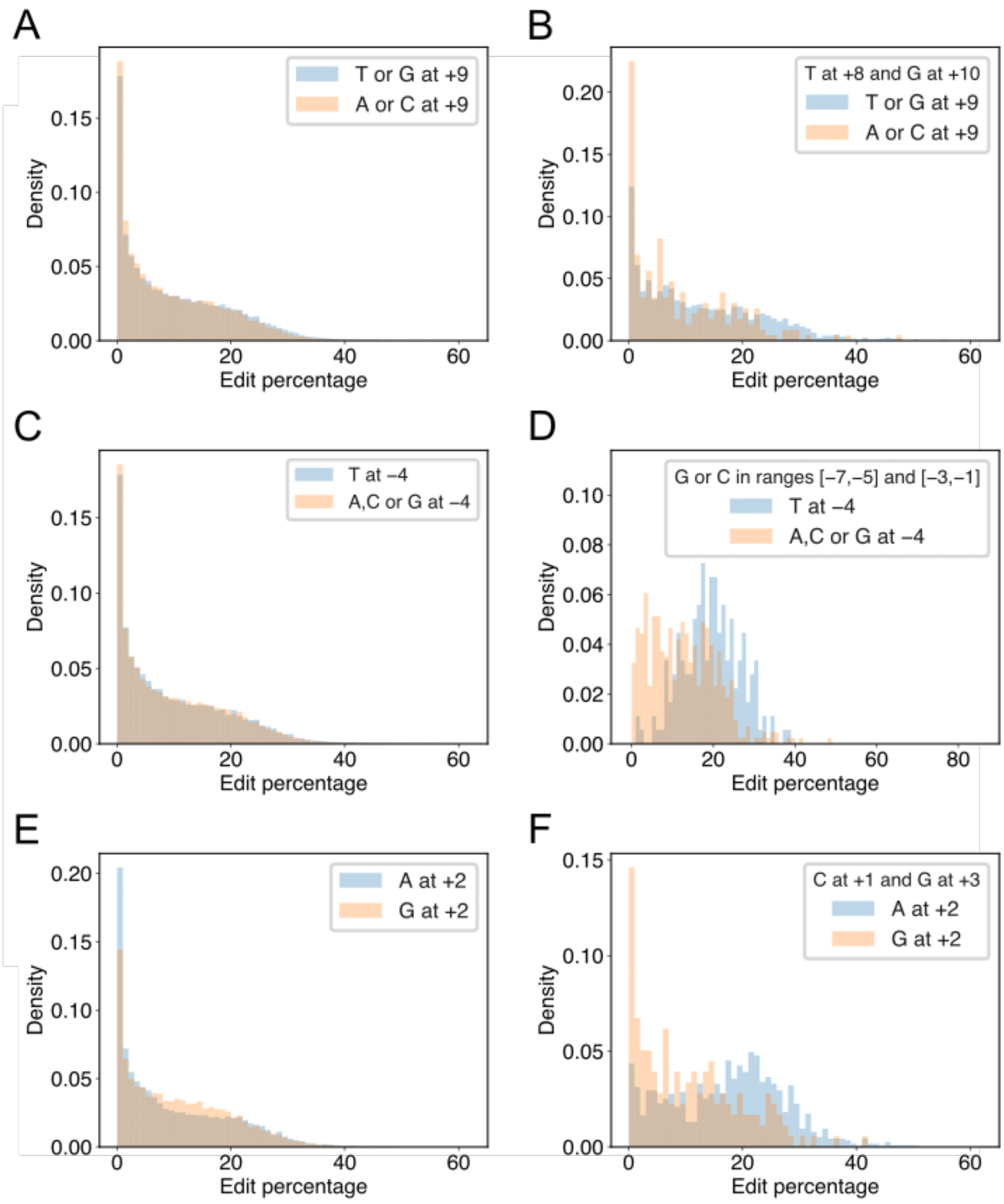


**Figure S8:** Edit percentages at target sites grouped into sequence categories. (A) Histograms of edit percentages at target sites with T or G at +9 (blue) vs. A or C at +9 (orange) relative to the nick site. (B) Same as in (A), but target sites are conditioned to have T at +8 and G at +10. (C) Histograms of edit percentages at target sites with T at -4 (blue) vs. A, C, G at -4 (orange). (D) Same as in (C), but target sites are conditioned to have G or C in the remaining positions of [-7,-1]. (E) Histograms of edit percentages at target sites with A at +2 (blue) vs. G at +2 (orange). (F) Same as in (E), but target sites are conditioned to have C at +1 and G at +3.

### SUPPLEMENTARY TABLES

| Gene | Protospacer | RT template | PBS |
|---|---|---|---|
| **ABCD1** | CTCGCGTGTGGTGGCCAACT | GAAGGCGATCTCCTTCGAGT | TGGCCACCACACG |
| **CACNA1A** | GTCGTAGCACCCGGAGGACT | ACGATTAAATCCCTCTGAGT | CCTCCGGGTGCTA |
| **CDKN2A** | GCATGGAGCCTTCGGCTGAC | GCGGCCGTGGCCAGCTAGTC | AGCCGAAGGCTCC |
| **DES** | CCCGCCGAAGGTGCGGCGGT | CCAGCGCGTGTCCTTCTACC | GCCGCACCTTCGG |
| **DLL3** | CGCTGCCGCGCCGGCTTCGC | GCTCGCAGCGAGGATCCGCG | AAGCCGGCGCGGC |
| **GRM6** | GAGCGCGTCGTGGCCGTCGT | AGCTGGCCGAGGCGCTCACG | ACGGCCACGACGC |
| **PURA** | TCTCTCCATGTCAGTGGCCG | GTAGTCGCGGAACTACACGG | CCACTGACATGGA |
| **SLC6A8** | ATTTCATCCAGGGGCAAGGT | TGTATGATCGGGTACTGACC | TTGCCCCTGGATG |
| **TERT** | AGTGCCTGGTGTGCGTGCCC | GGCGGCCGTGCGTCTCAGGG | CACGCACACCAGG |
| **TRIO** | CGGGCAACACCCTGCGCAAG | ACGGGGCTGGTGAGTCACTT | CAACACCCTGCGC |
| **ABCC6** | TGCCCCGCGCAGGGCTCAGC | CGCCGATGGCCGCGCATGCT | GAGCCCTGCGCGG |
| **ANKRD11** | CGATGGCGGCCAGCGTCTGC | ACGCGGGAGGTGATCTAGCA | GACGCTGGCCGCC |
| **ATM** | TATATATATTCTCTATTTAA | CTTTTTTTGCACCTACTTTA | AATAGAGAATATA |
| **CHRNE** | CTAAGCCCCGCCCCTGCCCC | CAGCTCCAGGAGAACGTGGG | GCAGGGGCGGGGC |
| **CREBBP** | GGGGTGGGGGGGCCGGCACC | GGGAAGCCTACCAGCTAGGT | GCCGGCCCCCCCA |
| **CYP11B1** | GGAAGGAGCACTTTGAGGCC | TGGAAGATGCAGTCCTAGGC | CTCAAAGTGCTCC |
| **CYP21A2** | CTGCGGTGCCCGCGTGTGCC | CGCCAGCGGCTCGCTCAGGC | ACACGCGGGCACC |
| **HEXB** | CCAAGCCGGGGCCGGCGCTG | AAGAGCGGCAGGGGCTACAG | CGCCGGCCCCGGC |
| **INS** | CCTCTGCCTCGCCGCTGTTC | GCGCAGAGCAGGTTCTGGAA | CAGCGGCGAGGCA |
| **LMNA** | CGCTGCCAACCTGCCGGCCA | CTGGGACGGGGTCTCTATGG | CCGGCAGGTTGGC |
| **PKD1** | GGACACTGCTGGCTCCACAC | CTCCCTGCTGGGCCCGTGTG | TGGAGCCAGCAGT |
| **RAX2** | GCCAGGCCTCCCCCGCCTCC | CCCCGGGCCCGGGCGCAGGA | GGCGGGGGAGGCC |
| **SYNJ1** | TGCGGAAGAGATGGGCCTGC | GCGTCACTTCCGCTTCAGCA | GGCCCATCTCTTC |
| **UMOD** | ACGATGCCCTCGTCGCTGGA | TCAATGGCACGCATCTGTCC | CGGATGCGTGCCA |
| **VHL** | CCCGTATGGCTCAACTTCGA | AGGGCTGCGGCTCGTCGTCG | AAGTTGAGCCATA |

**Table S1:** PegRNA designs used in the in-house experiments

| Gene | Forward | Reverse |
|---|---|---|
| **ABCD1** | GGCTGTGACTTCCTACACCC | TCAAACCGCTAGGATCGCAG |
| **CACNA1A** | AGACTTCTAGGCCTGGGAGG | CTGCTTCCCAGATCACGGTT |
| **CDKN2A** | GATTTGAGGGACAGGGTCGG | ACCCTCTACCCACCTGGATC |
| **DES** | CTCGCCGCATCCACTCTC | GAAGCGGTCATTGAGCTCCT |
| **DLL3** | GGAAGGCCAGAGGGTTCAAA | CACACGTGCCGCCGTTAG |
| **GRM6** | GAAGAAGGAGCAGGGCGTG | TTTGGGTGGAGCCGATTCAG |
| **PURA** | ACAAGCGCTTCTACCTGGAC | GTCGATGAGCTTGGCCAGAG |
| **SLC6A8** | GACTGATGTGAGTGGGGTGG | TAGACCAGCGGCTCGTAGTA |
| **TERT** | CGAGTTTCAGGCAGCGCT | CAGCTCCTTCAGGCAGGAC |
| **TRIO** | CCACTGGCTTCCCTGCAG | GGCAGAGCCTCACCTCCT |
| **ABCC6** | CCGAGCAGTCTGCCCAGAGACT | CTGGCAGTTTGCAACCCCCACC |
| **ANKRD11** | TCCCTCAGCGCATGACCAGGAA | AGAGAAGGCAGTGGCTCTCCCG |
| **ATM** | AGGTGTGCTTCCTCCCCCAACA | TCATCCCCCTGCAACTCAGAATGT |
| **CHRNE** | GGAGGAGGAACGGTGATGTC | CACCGGGGTGGGCCTTAG |
| **CREBBP** | TTCTCCTACCTCAGCACCGCCC | GTTGGGTCGGGGCACATTCAGG |
| **CYP11B1** | CCCCAGTTCTGCCAGCCTGAAC | CCGACACCCAAGTCTCCCTGCT |
| **CYP21A2** | GGGCCTGCCGTGAAAATGTGGT | AACTGAGGTACCCGGCTGGCAT |
| **HEXB** | TCTGACTCGGTGACTCACCCGC | CGAAACGCTTCCTCCAGCAGGG |
| **INS** | GGCCTGTAGGTCCACACCCAGT | ATCTCTCTCGGTGCAGGAGGCG |
| **LMNA** | AGCCAACCCAGATCCCGAGGTC | CAGACTCGGTGATGCGAAGGCG |
| **PKD1** | GAGGTGCCTTGTGTGGTCGGTG | GCGGCACCTGTGATGTTGAGGA |
| **RAX2** | TGCAGAGGGGGTCCGAGGATTG | AGCCTGTGCATGTTCCTTGGCC |
| **SYNJ1** | GCCCACGCAAAAGGACCAAGGA | AGACTGGTCTTGGAGGCGTCCG |
| **UMOD** | GCACCCTGGACGAGTACTG | GCACCCTGGACGAGTACTG |
| **VHL** | TGAAGAAGACGGCGGGGAGGAG | CCCCGTCTGCAAAATGGACCCC |

**Table S2:** Primer pairs used to generate the DNA amplicons for next-generation sequencing and measure edit percentages.

| Predictive variable | Regression coefficient | *p*-value |
|---|---|---|
| Intercept | -1.59 | $1.87\times10^{-4}$ |
| $Log_{10}$ Distance to the nearest H3K9me3 peak | 0.48 | $2.52\times10^{-10}$ |
| $Log_{10}$ Distance to the nearest H3K27me3 peak | -0.04 | $2.52\times10^{-1}$ |

**Table S3**: Regression coefficients of the logistic regression model trained on endogenous target sites from Kim *et al.*'s data (1).

| Test set | RMSE= $\sqrt{\underset{\substack{10\le i\le 20\\ gene\in S}}{ave}\left(\Delta EP_{gene}(i)-\Delta\widehat{EP}_{gene}(i)\right)^2}$ | | Pearson correlation | |
|---|---|---|---|---|
| | $S$ = Training set | $S$ = Test set | Training set | Test set |
| EMX1 | 1.9 % | 4.2 % | 0.83 | 0.47 |
| HEK4 | 2.3 % | 2.2 % | 0.80 | 0.51 |
| RNF2 | 2.3 % | 2.4 % | 0.80 | 0.61 |
| FANCF | 2.3 % | 2.1 % | 0.79 | 0.49 |
| HEK3 | 2.2 % | 2.5 % | 0.77 | 0.79 |
| VEGFA | 2.3 % | 2.2 % | 0.78 | 0.80 |
| DNMT1 | 2.3 % | 1.8 % | 0.77 | 0.88 |
| RUNX1 | 2.2 % | 2.7 % | 0.80 | 0.62 |

**Table S4**: The 8-fold cross-validation prediction metrics of the OLS linear regression model trained on Anzalone *et al.*'s data (2). $\Delta EP$ is the observed stepwise difference in edit percentages and $\Delta\widehat{EP}$ is the predicted stepwise differences in edit percentages.

| Predictive variable | Regression coefficient | *p*-value |
|---|---|---|
| Intercept | -0.74 | $2.41\times10^{-1}$ |
| Penultimate Templated: A | -3.04 | $3.41\times10^{-5}$ |
| Penultimate Templated: C | -3.59 | $5.94\times10^{-6}$ |
| Penultimate Templated: T | -4.64 | $1.01\times10^{-6}$ |
| Last Templated: A | 3.27 | $1.05\times10^{-5}$ |
| Last Templated: C | 6.25 | $4.49\times10^{-12}$ |
| Last Templated: T | 6.51 | $3.26\times10^{-11}$ |

**Table S5**: Regression coefficients of the OLS linear regression model trained on Anzalone *et al.*'s data (2) to predict stepwise differences in edit percentages.

| Test set ID | Range of RTTL | RMSE= $\sqrt{\underset{\substack{10\le i\le 20\\ gene\in S}}{ave}\left(\Delta EP_{gene}(i)-\Delta\widehat{EP}_{gene}(i)\right)^2}$ | | Pearson correlation | |
|---|---|---|---|---|---|
| | | $S$ = Training set | $S$ = Test set | Training set | Test set |
| 0 | $10 \le i \le 12$ | 5.2% | 5.4% | 0.35 | 0.17 |
| 1 | | 5.2% | 5.0% | 0.33 | 0.41 |
| 2 | | 5.1% | 5.6% | 0.37 | 0.12 |
| 3 | | 5.1% | 5.5% | 0.32 | 0.46 |
| 4 | | 5.1% | 5.5% | 0.33 | 0.40 |
| 5 | | 5.2% | 4.9% | 0.33 | 0.45 |
| 6 | | 5.2% | 5.1% | 0.35 | 0.24 |
| 7 | | 5.2% | 5.0% | 0.34 | 0.35 |
| 8 | | 5.2% | 4.7% | 0.35 | 0.22 |
| 9 | | 5.2% | 5.2% | 0.32 | 0.47 |
| **Average** | | **5.2%** | **5.2%** | **0.34** | **0.33** |
| 0 | $12 \le i \le 15$ | 5.2% | 5.8% | 0.27 | 0.17 |
| 1 | | 5.2% | 5.7% | 0.26 | 0.27 |
| 2 | | 5.2% | 5.5% | 0.25 | 0.29 |
| 3 | | 5.3% | 5.1% | 0.26 | 0.26 |
| 4 | | 5.2% | 5.8% | 0.26 | 0.29 |
| 5 | | 5.3% | 4.9% | 0.26 | 0.24 |
| 6 | | 5.2% | 5.4% | 0.26 | 0.22 |
| 7 | | 5.3% | 4.6% | 0.25 | 0.34 |
| 8 | | 5.3% | 4.4% | 0.26 | 0.26 |
| 9 | | 5.3% | 5.0% | 0.27 | 0.16 |
| **Average** | | **5.3%** | **5.2%** | **0.26** | **0.25** |
| 0 | $15 \le i \le 20$ | 5.0% | 5.2% | 0.34 | 0.30 |
| 1 | | 5.0% | 5.2% | 0.34 | 0.30 |
| 2 | | 5.0% | 4.7% | 0.33 | 0.32 |
| 3 | | 4.9% | 5.9% | 0.34 | 0.28 |
| 4 | | 5.0% | 5.0% | 0.33 | 0.36 |
| 5 | | 5.0% | 4.6% | 0.33 | 0.32 |
| 6 | | 5.0% | 5.1% | 0.32 | 0.48 |
| 7 | | 5.0% | 5.4% | 0.33 | 0.36 |
| 8 | | 5.0% | 4.7% | 0.34 | 0.28 |
| 9 | | 5.1% | 4.1% | 0.33 | 0.31 |
| **Average** | | **5.0%** | **5.0%** | **0.33** | **0.33** |

**Table S6:** The 10-fold cross validation prediction metrics of the OLS linear regression model trained on integrated target sites from Kim *et al*. (1). $\Delta EP$ is the observed stepwise difference in edit percentage and $\Delta\widehat{EP}$ is the predicted stepwise difference.

| **Predictive variable** | **Regression coefficient** | ***p*-value** |
|---|---:|---|
| Intercept | -3.76 | $<10^{-300}$ |
| Penultimate Templated: A | 3.62 | $<10^{-300}$ |
| Penultimate Templated: C | 1.61 | $2.13\times10^{-9}$ |
| Penultimate Templated: T | 2.40 | $<10^{-300}$ |
| Last Templated: A | -0.10 | $7.10\times10^{-1}$ |
| Last Templated: C | 1.08 | $4.18\times10^{-5}$ |
| Last Templated: T | 0.15 | $5.92\times10^{-1}$ |

**Table S7**: Regression coefficients of the OLS linear regression model trained on integrated target sites from Kim *et al*. (1) to predict stepwise differences in edit percentages for RTTL in the range [10,12].

| **Predictive variable** | **Regression coefficient** | ***p*-value** |
|---|---:|---|
| Intercept | -3.62 | $<10^{-300}$ |
| Penultimate Templated: A | -1.63 | $1.86\times10^{-10}$ |
| Penultimate Templated: C | -1.43 | $1.59\times10^{-8}$ |
| Penultimate Templated: T | -1.55 | $1.88\times10^{-9}$ |
| Last Templated: A | 2.31 | $<10^{-300}$ |
| Last Templated: C | 2.43 | $<10^{-300}$ |
| Last Templated: T | 2.52 | $<10^{-300}$ |

**Table S8**: Regression coefficients of the OLS linear regression model trained on integrated target sites from Kim *et al*. (1) to predict stepwise differences in edit percentages for RTTL in the range [12,15].

| **Predictive variable** | **Regression coefficient** | ***p*-value** |
|---|---:|---|
| Intercept | -1.76 | $2.66\times10^{-4}$ |
| Penultimate Templated: A | -2.63 | $<10^{-300}$ |
| Penultimate Templated: C | -2.81 | $<10^{-300}$ |
| Penultimate Templated: T | -2.87 | $<10^{-300}$ |
| Last Templated: A | 2.04 | $<10^{-300}$ |
| Last Templated: C | 2.91 | $<10^{-300}$ |
| Last Templated: T | 2.45 | $<10^{-300}$ |

**Table S9**: Regression coefficients of the OLS linear regression model trained on integrated target sites from Kim *et al*. (1) to predict stepwise differences in edit percentages for RTTL in the range [15,20].

| RTTL / PBSL | 10 | | 12 | | 15 | | 20 | |
|---|---|---|---|---|---|---|---|---|
| | ***r*** | ***p*-value** | ***r*** | ***p*-value** | ***r*** | ***p*-value** | ***r*** | ***p*-value** |
| **7** | 0.70 | $8.98{\times}10^{-266}$ | 0.69 | $1.60{\times}10^{-255}$ | 0.67 | $1.03{\times}10^{-232}$ | 0.64 | $9.37{\times}10^{-211}$ |
| **9** | 0.71 | $3.01{\times}10^{-280}$ | 0.68 | $1.57{\times}10^{-250}$ | 0.69 | $4.19{\times}10^{-259}$ | 0.68 | $1.23{\times}10^{-248}$ |
| **11** | 0.68 | $5.88{\times}10^{-242}$ | 0.65 | $3.18{\times}10^{-221}$ | 0.64 | $7.77{\times}10^{-212}$ | 0.67 | $9.59{\times}10^{-232}$ |
| **13** | 0.68 | $2.73{\times}10^{-241}$ | 0.66 | $1.92{\times}10^{-222}$ | 0.62 | $5.44{\times}10^{-192}$ | 0.66 | $2.71{\times}10^{-218}$ |
| **15** | 0.67 | $6.81{\times}10^{-234}$ | 0.62 | $1.64{\times}10^{-193}$ | 0.62 | $1.02{\times}10^{-195}$ | 0.65 | $2.06{\times}10^{-211}$ |
| **17** | 0.63 | $7.05{\times}10^{-192}$ | 0.62 | $1.66{\times}10^{-190}$ | 0.63 | $5.60{\times}10^{-197}$ | 0.60 | $2.11{\times}10^{-160}$ |

**Table S10**: Pearson correlation coefficient *r* between the observed edit percentages and predicted edit percentages of the elastic net model trained on integrated target sites from Kim *et al*. for each pair of PBSL and RTTL (1).

| $q_V$ / $q_A$ | 1.00 | | 1.20 | | 1.40 | | 1.60 | |
|---|---|---|---|---|---|---|---|---|
| | $ave(EP_{max})$ | $sd(EP_{max})$ | $ave(EP_{max})$ | $sd(EP_{max})$ | $ave(EP_{max})$ | $sd(EP_{max})$ | $ave(EP_{max})$ | $sd(EP_{max})$ |
| **1.00** | 50.91 | 0.05 | 50.97 | 0.00 | 50.63 | 0.43 | 50.55 | 0.16 |
| **1.20** | 50.77 | 0.09 | 50.97 | 0.00 | 50.91 | 0.09 | 50.62 | 0.45 |
| **1.40** | 50.45 | 0.14 | 50.97 | 0.00 | 50.97 | 0.00 | 50.82 | 0.22 |
| **1.60** | 50.09 | 0.22 | 50.97 | 0.00 | 50.97 | 0.00 | 50.95 | 0.06 |

**Table S11**: Approximations of the maximum DNN-predicted edit percentage for PBSL=13 and RTTL=15 by modified simulated annealing using the indicated coarse grid values of $q_A$ and $q_V$. For each combination of $q_A$ and $q_V$, we initialized 10 independent Markov chains with a fixed set of 10 random sequences and simulated them for $2 \times 10^6$ iterations; the average and standard deviation of the maximum edit percentages $EP_{max}$ found by the 10 chains are shown.

| $q_V$ / $q_A$ | 1.00 | | 1.20 | | 1.40 | | 1.60 | |
|---|---|---|---|---|---|---|---|---|
| | $ave(i_{max})$ | $sd(i_{max})$ | $ave(i_{max})$ | $sd(i_{max})$ | $ave(i_{max})$ | $sd(i_{max})$ | $ave(i_{max})$ | $sd(i_{max})$ |
| **1.00** | 1,266,690 | 619,877 | 21,563 | 11,652 | 2,278 | 849 | 1,154 | 705 |
| **1.20** | 1,576,217 | 372,842 | 27,119 | 11,116 | 15,099 | 18,430 | 1,525 | 747 |
| **1.40** | 1,432,015 | 427,860 | 75,923 | 29,110 | 6,916 | 2,829 | 2,362 | 1,679 |
| **1.60** | 1,066,327 | 666,246 | 169,532 | 81,692 | 12,004 | 7,082 | 127,544 | 127,544 |

**Table S12**: Number of iterations taken by modified simulated annealing to find an optimal sequence approximately maximizing the DNN-predicted edit percentage for PBSL=13 and RTTL=15. For each indicated combination of $q_A$ and $q_V$, we examined the 10 independent Markov chains described in Table S11 and recorded their respective iteration number $i_{max}$ corresponding to the first instance of the maximum in each chain; the average and standard deviation of $i_{max}$ were then calculated across the 10 chains.

| $q_V$ / $q_A$ | 1.25 | | 1.30 | | 1.35 | |
|---|---|---|---|---|---|---|
| | $ave(EP_{max})$ | $sd(EP_{max})$ | $ave(EP_{max})$ | $sd(EP_{max})$ | $ave(EP_{max})$ | $sd(EP_{max})$ |
| **1.05** | 50.96 | 0.04 | 50.95 | 0.06 | 50.91 | 0.09 |
| **1.10** | 50.97 | 0.00 | 50.97 | 0.00 | 50.90 | 0.12 |
| **1.15** | 50.97 | 0.00 | 50.97 | 0.00 | 50.97 | 0.00 |
| **1.20** | 50.97 | 0.00 | 50.97 | 0.00 | 50.97 | 0.00 |

**Table S13**: Approximations of the maximum DNN-predicted edit percentage for PBSL=13 and RTTL=15 by modified simulated annealing using the indicated refined grid values of $q_A$ and $q_V$. For each combination of $q_A$ and $q_V$ at a step size of 0.05 in the ranges [1.05,1.20] and [1.25,1.35], respectively, we initialized 10 independent Markov chains with the same fixed set of 10 random sequences from Table S11 and simulated them for $2 \times 10^6$ iterations; the average and standard deviation of the maximum edit percentages $EP_{max}$ found by the 10 chains are shown.

| $q_V$ / $q_A$ | 1.25 | | 1.30 | | 1.35 | |
|---|---|---|---|---|---|---|
| | $ave(i_{max})$ | $sd(i_{max})$ | $ave(i_{max})$ | $sd(i_{max})$ | $ave(i_{max})$ | $sd(i_{max})$ |
| **1.05** | 37,811 | 71,193 | 6,163 | 2,729 | 9,296 | 9,814 |
| **1.10** | 27,774 | 31,583 | 11,287 | 9,588 | 6,485 | 5,485 |
| **1.15** | 12,192 | 8,039 | 10,674 | 8,115 | 21,497 | 47,803 |
| **1.20** | 23,179 | 12,947 | 12,793 | 15,067 | 19,757 | 39,189 |

**Table S14**: Number of iterations taken by modified simulated annealing to find an optimal sequence approximately maximizing the DNN-predicted edit percentage for PBSL=13 and RTTL=15. For each indicated combination of $q_A$ and $q_V$, we examined the 10 independent Markov chains described in Table S13 and recorded their respective iteration number $i_{max}$ corresponding to the first instance of the maximum in each chain; the average and standard deviation of $i_{max}$ were then calculated across the 10 chains.
